# Cathelicidin senses enteric pathogen *Salmonella typhimurium/*LPS for colonic chemokine generation: a new innate immune role for a host defense peptide

**DOI:** 10.1101/389726

**Authors:** Ravi Holani, Fernando Lopes, Graham A. D. Blyth, Humberto Jijon, Derek M. McKay, Morley D. Hollenberg, Eduardo R. Cobo

## Abstract

The mechanisms by which epithelia identify and respond to pathogens are manifold, nuanced and complex. Here, using human-colon derived HT29 epithelial cells, mouse and human primary colonoids, and cathelicidin null *(Cramp)* mice, we report a novel immunoregulatory role for the antimicrobial peptide, cathelicidin, that was found to recognize and synergise with *Salmonella typhimurium* or its derived virulence factor lipopolysaccharide (LPS) to promote epithelial synthesis of the chemokine IL-8/KC for neutrophil recruitment/activation during infectious colitis. Mechanistically, cathelicidin facilitated the internalization of LPS via GM1 lipid rafts and subsequent TLR4 activation to promote IL-8 production. Furthermore, IL-8 output required the integrated activity of two signal transduction pathways: NF-κB and MEK 1/2 kinase signaling was required for IL-8 mRNA synthesis, while Src-EGFR-p38MAPK (NF-κB independent) activity underlay IL-8 mRNA stabilization. This immunomodulatory function of cathelicidin was key in colon defense, since *Cramp^−/−^* mice infected with a natural murine Gram negative intestinal pathogen, *Citrobacter rodentium,* displayed diminished KC secretion, impaired mobilization and reduced clearance of the bacteria. Occurring at concentrations lower than those necessary for anti-microbial activity, cathelicidin’s capacity to sense pathogens/LPS and enhance neutrophil recruitment reveals a novel function for this peptide in directing innate immunity which may be of pivotal importance in the control of infections colitis.

**Author summary:** The gut lining has a well regulated immune system that tolerates resident bacteria and does not respond to them. However, when pathogenic bacteria enter, there needs to be a protective response. How the gut lining ‘switches’ from passive to protective is of interest. In our study, we determined host defense cathelicidin peptide (either naturally occurring or administered) “instructs” the colon lining to produce a compound (IL-8) that attracts white blood cells in response to a pathogen *(Salmonella typhimurium)* or lipopolysaccharide, a component of this pathogen’s cell wall. We discovered a novel mechanism by which cathelicidin facilitates uptake of lipopolysaccharide by the lining of the colon and how it activates receptors to increase synthesis and release of IL-8. In addition, we also detected a synergistic action between cathelicidin and intestinal pathogens in laboratory cultures of colon tissues from mice and humans, as well as in a mouse model of colitis with another pathogenic bacterium. Cathelicidin induced production of IL-8 which attracted and stimulated more white blood cells. Therefore, in addition to potential direct actions to supress harmful bacteria, cathelicidin also acts as a biological sensor in the gut lining, recognizing pathogens or factors they produce and increasing white cell responses.

## Introduction

Infectious colitis caused by Gram-negative enteric pathogens, such as *Salmonella typhimurium,* is a common cause of diarrhoea and a leading cause of morbidity and mortality among young children, particulary in developing countries (1). Neutrophils migrating to the colonic site of inflammation constitute an early innate immune defense in *S. typhimurium-* induced acute colitis (2, 3). Indeed, mice depleted of neutrophils (anti-Gr-1 antibodies) and infected with *S. typhimurium* had higher bacterial loads at extra-intestinal sites, particularly in mesenteric lymph nodes, spleen, and liver (4).

Mechanistically, directional movement of neutrophils into the intestinal mucosa is largely regulated by the C-X-C motif-containing chemokine ligand 8 (CXCL8; or interleukin (IL)-8). IL-8 is expressed by both haematopoietic (neutrophils and macrophages) and non-haematopoietic (colonic and skin epithelia) cells (5). In mice, CXCL1-chemokine (C-X-C motif) ligand 1, also termed keratinocyte derived chemokine (KC), is the homologue of IL-8, with similar functions in regulating neutrophil recruitment (6). Mice deficient in KC and challenged with the abiotic colitis inducer dextran sodium sulphate (DSS) have reduced neutrophil recruitment, accompanied by severe bloody diarrhoea and a loss of intestinal barrier integrity (7). At the cellular level, expression of IL-8/KC by intestinal epithelium can be induced by virulence factors from Gram-negative bacteria, including the major component of the outer bacterial membrane, lipopolysaccharide (LPS) (8). This balance between *S. typhimurium* invasion and influx of neutrophils due to IL-8 expression is central in pathogenesis of infectious colitis (9, 10). Mechanisms regulating IL-8 secretion in the intestine that can increase its production may stimulate an influx of neutrophils, thereby enhancing protection during infectious colitis.

Colonic epithelium and neutrophils are sources of cathelicidin, a naturally occurring, evolutionarily conserved small cationic amphiphilic peptide (11). Structurally, cathelicidin has an N-terminal domain containing a signal peptide, a conserved central cathelin domain, and a variable C-terminal domain with antimicrobial and immunomodulatory properties (12). There is only one type of cathelicidin in humans (cathelicidin antimicrobial peptide, hCAP-18, which yields a C-terminal active fragment termed leucine-leucine, a peptide with 37 amino acid residues, LL-37), whereas in mice the homologous peptide is termed cathelicidin-related-antimicrobial-peptide, Cramp (13). In addition to antimicrobial actions, cathelicidins have immunomodulatory effects. For instance, cathelicidin reduces production of pro-inflammatory cytokines TNF-α, IL-1β, IL-6 and IL-8 in cultured (THP-1) monocytes (14). Cathelicidin secretion by intestinal epithelial cells can be induced by either direct contact with pathogens or with bacterial by-products such as butyrate or LPS (15, 16). However, underlying mechanism(s) whereby cathelicidin modulates expression of neutrophil chemoattractants, including IL-8, in the presence of enteric pathogens, and resulting impacts on neutrophil recruitment and colitis, remain unknown.

In this study, we hypothesized that the invading pathogen and cathelicidin cooperate to promote synthesis of IL-8 (or KC) in the colonic epithelium. We therefore evaluated the process whereby LL-37, in association with either *S. typhimurium* or its derived LPS, causes transcriptional upregulation of *IL-8* (or KC) mRNA and its secretion in cultured human colonic epithelial cells, colonic organoids derived from humans and mice, and cathelicidin deficient Cramp-null mice. This induced IL-8 biosythesis was facilitated by sequential crosstalk among toll-like receptor 4 (TLR4), epidermal growth factor receptor (EGFR) kinase, sarcoma family (Src) kinases, p38 mitogen-activated protein kinase (p38MAPK), MAPK kinase (MEK1/2) and nuclear factor kappa-light-chain-enhancer of activated B cells (NF-κB) in colonic epithelium. The importance of cathelicidin for recruitment and activation of colonic neutrophils in response to Gram negative pathogens was further confirmed in a murine model of *Citrobacter rodentium* colitis using both wild type *(Cramp^+/+^)* and cathelicidin deficient Cramp-null mice.

## Materials and methods

### Colonic epithelial cell culture

Human colon epithelial adenocarcinoma-derived HT29 and T84 cells (originally obtained from ATCC and kindly provided by Dr. K. Chadee, University of Calgary) were cultured in Dulbecco’s Modified Eagle’s Media (DMEM; Gibson, Life Technologies) containing 4.5 g/L glucose with 10% fetal bovine serum (FBS; Benchmark Gemini BioProducts), 1 mM sodium pyruvate (Gibco, Life Technologies), and 1% penicillin (100 U ml^-1^)/ streptomycin (100 μg ml^-1^; HyClone Thermo, Fisher Scientific) in humidified environment with 5% CO_2_. T84 cells were grown in transwell chambers to form polarized monolayers with a transepithelial electrical resistance (TER) of >1000 Ω/cm^2^ (monitored by an electro-voltameter; World Precision Instruments). For experiments, cells were cultured in DMEM without FBS or antibiotics.

### Signal transduction reagent probes

Signal pathway involvement was investigated by pharmacologically blocking human epidermal growth factor receptor 2 (EGFR 2/ErbB2; TAK 165, 366017-09-6; Tocris Bioscience), formyl peptide receptor-like 1(FPRL1; WRW4, 2262; Tocris Bioscience), matrix metalloproteinases (MMP; GM6001, 2983; Tocris Bioscience), P2X purinoceptor 7 (P2X7; A740003, 3701; Tocris Bioscience), EGFR-kinase (AG1478 hydrochloride, 1276; Tocris Bioscience), sarcoma family kinases (Src-kinases; PP1, Calbiochem) and mitogen-activated protein kinase (MAPK) kinase (MEK, also known as MAPKK) 1/2 (PD98059, 9900; Cell Signaling Technology and U0126, 1144; Tocris Bioscience), p38 MAPK (SB203580, 1202; Tocris Bioscience), extracellular signal-regulated kinases (ERK; FR180204, 3706; Tocris Bioscience), and nuclear factor kappa-light-chain-enhancer of activated B cells (NF-κB) activating kinase, i.e. IκB kinase β (IKKβ; PS-1145; Cayman Chemical). Endocytic processes were investigated by inhibiting endocytosis (D15, 2334; Tocris Bioscience), actin polymerization (Cytochalasin D, C8273; Sigma-Aldrich), and lipid raft formation (Mevinolin, M2147, HMG-CoA reductase inhibitor; Sigma-Aldrich and methyl-β-cyclodextrin, 332615, cholesterol solubilizing agent; Sigma-Aldrich). Cycloheximide (2112; Cell Signaling Technology) and actinomycin D (15021; Cell Signaling Technology) were used to assess gene translation and transcription, respectively. The concentrations used for the above mentioned inhibitors were either based on their IC_50_ values as recommended by manufacturer(s) or obtained from previous studies (14, 17–20).

### Murine models

All animal experiments using male 8-wk-old *Cramp^+/+^* and *Cramp^−/−^* C57BL/6 mice (B6.129X1-*Camp^tm1Rlg^*/J; The Jackson Laboratory) housed in a pathogen-free environment (Health Sciences Animal Resources Centre, University of Calgary) were conducted in accordance with Canadian Guidelines for Animal Welfare (CGAW).

For systemic LPS challenge, *Cramp^−/−^* mice were injected intraperitoneally (ip) with LPS derived from *S. typhimurium* (L6143; Sigma-Aldrich) ± synthethic LL-37 peptide (>98.6% purity, H-6224.0005; Bachem) (both at 1 μg/g), diluted in pyrogen-free saline. Control mice were sham injected with saline. Mice were humanely euthanized 3 h post challenge using 5% isoflurane anaesthesia, followed by cervical dislocation. Sections of distal colon were sampled, weighed and suspended in phosphate buffered saline (PBS, Gibco, Life Technologies; 1x) containing protease inhibitors (87786; ThermoFisher Scientific), at a final concentration of 50 mg tissue wet-weight/mL (21). Tissues were homogenized using an Omni TH-tissue homogenizer (CA11005-598; VWR), centrifuged (300 x g, 5 min, 4°C) and KC cytokine quantified using DuoSet ELISA kit (DY453; R&D Systems) in 100-μL aliquots of homogenized tissue, according to manufacturer’s recommendations. Data were represented as absolute values in pg/mL, normalized to tissue weight.

For the *C. rodentium* colitis model (22), the *C. rodentium* strain DBS-100 (kindly provided by Dr. A Buret, University of Calgary) was cultured on McConkey agar plates (16 h, 37°C) and single colonies sub-cultured in LB broth (5 mL, 16 h, 37°C) without shaking. The following day, an aliquot of the bacterial culture (1 mL) was transferred into a tissue culture flask containing fresh LB broth (50 mL) and cultured (4 h, 37°C) in a shaking incubator. Bacterial cells were centrifuged (3000 x g, 10 min, 4°C), washed (2 times) with sterile PBS (1x), and resuspended at ~ 5 x 10^8^ CFU/mL. Each mouse (wild-type *Cramp^+/+^* or cathelicidin-null *Cramp^−/−^*) was gavaged orally with *C. rodentium* (~ 1 x 10^8^ CFU in 200 μL), an adequate dose for establishing infection in C57BL/6 mice (23, 24). Control mice were sham-gavaged with 200 μL PBS (1x). Mice were humanely euthanized at day 7 post infection (pi), followed by collection of distal colon tissues. Tissues were weighed and suspended in either hexadecyltrimethylammonium bromide (HTAB) buffer (50 mg/mL) for myeloperoxidase (MPO) activity determination or in sterile-PBS containing protease inhibitor cocktail (50 mg/mL) for ELISA (21). Tissues were homogenized using a tissue homogenizer and centrifuged (300 x g, 5 min, 4°C). To determine MPO activity, tissue supernatant (7 μL) was mixed with an O-dianisidine solution (200 μL), containing O-dianisidine (0.5 mM) and hydrogen peroxide (1%) in potassium phosphate buffer. Optical absorbance density was recorded at 410 nm. MPO activity was determined in units (U)/mg and represented as fold increase in MPO activity with respect to wild type control mice. For KC cytokine quantification, ELISA was performed as discussed above.

To determine fecal shedding of *C. rodentium,* fresh fecal pellets (1 per mouse) were obtained aseptically from wild-type *Cramp^+/+^* and cathelicidin-null *Cramp^−/−^* at day 7 pi. These fecal pellets were then suspended in sterile PBS (50 mg/mL) and serial dilutions were plated on McConkey agar plates (24 h, 37°C). Bacterial colonies were then counted and data were represented as fold increase in CFU/g normalized to wild-type *Cramp^+/+^* infected mice. To confirm the identity of bacterial colonies on McConkey agar, a qPCR validation was done using *C. rodentium* specific primers against *espB* (extracellularly secreted protein B) gene (Forward primer: 5’-ATGCCGCAGATGAGACAGTTG-3’ and Reverse primer: 5’-CGTCAGCAGCCTTTTCAGCTA-3’) as described previously (25).

### Murine and human colonoids

Mini 3D-gut colonoids were developed from inducible pluripotent intestinal stem cells (iPSCs) isolated from murine colonic crypts and perpetuated in presence of specialized growth factor-enriched media (26). The full-length colon from male 8-wk-old wild-type mice was collected and transferred to PBS (1x). Fat was carefully removed and the colon sectioned into smaller pieces (< 1 mm) that were placed in crypt isolation buffer containing PBS (1x), ethylenediaminetetraacetic acid (EDTA; 2 mM; 6381-92-6; Sigma-Aldrich), sucrose (43.4 mM; S0389; Sigma-Aldrich), and dithiothreitol (DTT; 0.5mM; 3483-12-3; Sigma-Aldrich). After removing debris, tissue was repeatedly centrifuged (300 x g, 5 min) and seeded onto Matrigel matrix (354234; Corning) in a 12-well plate containing IntestiCult™ Organoid Growth Medium (06005; Stemcell Technologies) supplemented with antibiotic-antimycotic mixture (15240062; ThermoFisher Scientific). Medium was replaced every alternate day until day 10, when tissue acquired colonoid architecture (S1 Fig). At this point, cells were scraped from wells and seeded onto matrigel-coated 12 well-transwell chambers to yield a polarized 3D monolayer with the basolateral surface facing the underlying matrigel matrix. Culture medium was replaced in both apical and basolateral compartments on day 2, and on alternate days in basolateral compartment only until day 10 (26, 27).

Human colonic crypts were isolated as described above for mouse colonoids. Upon isolation, crypts were cultured on Matrigel matrix in a 24-well plate for 10 d until colonoid morphology was apparent (S1 Fig).

These colonoids were then treated with LPS and LL-37 or scrambled LL-37 (sLL-37; only in mouse colonoids), either alone or in combination, and supernatants collected for quantification of KC (as dicussed above) or IL-8, using DuoSet ELISA kit (DY208; R&D Systems) as per manufacturer’s recommendations. Data for KC cytokine were represented as absolute values in pg/mL. Data from human colonoids were normalized to LL-37 treatment group (IL-8 secretions in control and LPS treatment groups were below detection) and represented as fold increase in IL-8 secretion, to account for inter-patient variability.

#### Salmonella typhimurium

A *S. typhimurium* clinical strain LT2/ATCC 700720 (kindly provided by Dr. J. De Buck University of Calgary) was initially grown in Luria-Bertani (LB) Miller broth (IBI Scientific) in a shaker incubator (18 h, 37°C at 225 rpm) and streaked out on a LB agar plate. For experiments, five colonies were taken from agar plate and grown in LB broth (3 mL, 2 h, and 37 °C at 225 rpm). The resultant culture was serially 10 fold-diluted and dilutions streaked out on agar plate for 18 h, with the same dilutions subjected to OD600nm determination. A standard curve for colony forming units (CFU)/mL against OD_600nm_ was obtained. HT29 cells were challenged with *S. typhimurium* at multiplicity of infection (MOI) of 10:1 (2 x 10^7^ CFU/mL) for 4 h at 5% CO_2_ and 37 °C.

### Quantification of secreted cathelicidin

LL-37 in supernatants from HT29 cells infected with *S. typhimurium* was quantified using an ELISA kit (HK321; Hycult Biotechnology) as per manufacturer’s protocol. Data were represented as fold increase in LL-37 secretion normalized to control sham transfected group.

### Histological assessment and immunofluorescence in murine colon

Fresh colonic tissues (<1 cm) were fixed in 10% neutral buffered formalin for 3 h, transferred to ethanol 100% at 4°C, and then embedded in paraffin blocks. Sections (5 μm) were prepared and de-paraffinized by xylene substitute (Neo-Clear 65351-85 Millipore), followed by decreasing concentrations of ethanol and routine H&E staining. For immunofluorescence, after deparaffinization, slides were washed under running tap water (5 min), and rinsed in ice cold PBS with 0.05% Tween-20 (PBS-Tw; pH 7.2), followed by blocking in PBS-Tw containing 10% donkey serum (017-000-021; Jackson ImmunoResearch), 1% bovine serum albumin (BSA, 9048-46-8; Amresco), and 0.3 M glycine (1 h, RT). After rinsing with ice cold PBS-Tw, slides were incubated with primary Alexa647 tagged-anti-Lymphocyte antigen 6 complex locus G6D (Ly6G) antibody (5 μg/mL; MAB91671; R&D Systems) diluted in PBS (16 h, 4°C). Slides were rinsed with PBS-Tw, counterstained with 4’, 6-diamidino-2-phenylindole (DAPI, 62247; ThermoFisher Scientific) (1:1,000), and mounted with FluorSave (345789; Calbiochem). Slides were examined using wide-field immunofluorescence microscope (IX71; Olympus). Integrated fluorescence intensity per mouse was calculated using ImageJ 1.50i software (© National Institute of Health) in five randomly selected fields of view, normalized to their respective area, and averaged to obtain a single numerical value or mean fluorescence intensity (MFI). Data were reported as MFI from n= 4 mice.

### Colonic epithelial cell stimulation and detection of IL-8 transcription and secretion

HT29 and T84 cells were pre-treated (or not) for 1 h with signal pathway inhibitors listed above and maintained in culture medium without FBS or antibiotics during treatment. Cells were stimulated with *S. typhimurium,* its derived LPS (L6143, Sigma-Aldrich), Alexa Fluor^®^ 488-conjugated LPS from *S. minnesota* (L-23356; Molecular Probes), or lipid A from *S. minnesota* R595 (HC4057; Hycult Biotechnology). Effects of LPS on colonic cells was further studied using a TLR4-competitive inhibitor of LPS isolated from *Rhodobacter sphaeroides* (LPS: RS, tlrl-prslps; Invivogen) and polymyxin B sulfate salt (1405-20-5; Sigma-Aldrich). These treatments were combined with synthetic LL-37 amide trifluoroacetate salt or the LL-37 scrambled peptide (63708; Ana Spec). Cytotoxicity of LPS and LL-37 were determined using a Pierce™ LDH Cytotoxicity Assay Kit (88953; ThermoFisher Scientific) at various concentrations of LPS (10 ng/mL to 5 μg/mL) and LL-37 (1 to 30 μg/mL).

Secretion of human CXCL8 was quantified from supernatants from either HT29 or T84 cells, as described above. Secreted IL-8 was quantified as pg/mL/well, averaged over three wells per treatment per experiment (unless mentioned otherwise in figure legends), and represented as absolute values in pg/mL.

The LPS/LL-37 induced transcription of *IL-8* mRNA was quantified by quantitative real-time polymerase chain reaction (qPCR). HT29 cells were grown in 24-well plates and treated with LPS and LL-37, either alone or in combination, to trigger *IL-8* transcription. RNA was extracted using Ribozol^TM^ RNA extraction reagent (Amresco). For qPCR studies, 1 μg RNA was reverse-transcribed using qScript cDNA synthesis kit (Quantbio). RNA and DNA quality and quantity were assessed with a NanoVue Spectrophotometer (GE Healthcare Bio-Sciences Corp). Absence of contaminating genomic DNA from RNA preparations was verified using a minus-reverse transcriptase (RT) control (i.e., sample with all RT-PCR reagents except RT). qPCR was performed using a CFX-96 real time PCR system (BioRad). Each reaction mixture contained cDNA (100 ng), SsoAdvanced Universal SYBR Green Supermix (1x; BioRad), and 0.5 μM of each specific primer, in a final volume of 10 μL. Pre-designed primers (RT^2^ qPCR Primer Assay, Qiagen) specific for human *IL-8* (PPH00568A; NM_000584.3) and *glyceraldehyde-3-phosphate dehydrogensase (GAPDH)* (PPH00150F; NM_002046.5) were used. These primers were verified for specificity and efficiency to ensure amplification of a single product of the correct size with high PCR efficiency (> 95%), as indicated in MIQE guidelines (28). Reaction mixtures were incubated for 5 min at 95°C, followed by denaturation for 5 s at 95°C and combined annealing/extension for 10 s at 60°C, for a total of 40 cycles. *GAPDH,* along with two other housekeeping genes, *hypoxanthine-guanine phosphoribosyltransferase 1 (HGPRT1)* and *phosphoglycerate kinase 1 (PGK1)* were tested and determined invariable among treatment groups; therefore, *GAPDH* was used as a housekeeping gene for HT29 cells, as reported (29). Target gene mRNA values were corrected relative to the normaliser, *GAPDH.* Negative controls for cDNA synthesis and PCR procedures were included in all cases. Data were analysed using the 2^-ΔΔCT^ methods and results reported as mean fold change of target transcript levels in stimulated groups versus an untreated control group.

### Measuring mRNA stability (actinomycin D chase assay)

Stability of mRNA in HT29 cells in which *IL-8* was up-regulated by combined actions of LPS and cathelicidin was determined. Cells were washed (3 times) with PBS (1x) after a challenge with LPS ± LL-37 (for 2 h). Either actinomycin D alone or in combination with p38MAPK inhibitor (SB203580)/MEK1/2 inhibitor (PD98059) was added for the indicated intervals. Cells were washed with PBS (1x) and RNA was isolated using Ribozol™ RNA extraction reagent, as discussed above. Following cDNA synthesis and subsequent qPCR, data were represented as percentage *IL-8* mRNA remaining compared to actinomycin D only treatment as control.

### Western blot analysis of signal transduction pathway activation

Activation/phosphorylation of signal pathway components (p38MAPKinase, EGF receptor; and NF-κB) was monitored by western blot analysis. HT29 cell monolayers grown in 6-well plates were treated with LL-37 amide and LPS, either alone or in combination, rinsed with ice-cold PBS (pH 7.4) at variable intervals (as indicated in respective figures) and harvested by scraping into ice-cold PBS (1x; 1 mL). Upon centrifugation (1000 x g, 5 min, 4 °C), cell pellets were re-suspended in denaturing cell extraction buffer (DCEB; FNN0091; Thermo Fischer Scientific) and reconstituted with a protease inhibitor cocktail. Protein content in the solubilized cell sample was quantified using a Pierce bicinchoninic acid (BCA) colorimetric Protein Assay Kit (23225; Thermo Fisher Scientific). Extracted proteins were separated by SDS-polyacrylamide gel electrophoresis (7.5-15%, depending on molecular weight of protein) and transfered to methanol-activated polyvinylidene difluoride membranes (PVDF; 1620177; Bio-Rad). Membranes were blocked with 5% BSA (9048-46-8; Amresco) dissolved in Tris buffered saline plus 0.1% Tween 20 solution (TBST) for 2 h and probed (16 h, 4 °C) with the following specific primary antibodies (diluted in TBST 1:1,000): phospho-EGFR-Tyr1068 (2234; Cell Signaling Technology), EGFR (4267; Cell Signaling Technology), phospho-p38 MAPK-Thr180/Tyr182 (4511; Cell Signaling Technology), p38 MAPK (9212; Cell Signaling Technology), phospho-NF-κB p65-Ser536 (3033; Cell Signaling Technology), and human GAPDH-6C5 (1001; Calbiochem). Secondary antibodies used were horseradish-peroxidase-conjugate (HRP) goat anti-mouse IgG (H+L) (115-035-146; Jackson ImmunoResearch) and HRP goat anti-rabbit IgG (H+L) (115-035-144; Jackson ImmunoResearch), diluted in TBST (1:10,000), and incubated with membrane for 1 h at room temperature (RT). Blots were developed using the Clarity Western ECL Detection System (BioRad). Image capture and densitometric analyses were performed with the ChemiDoc MP Imaging system and ImageLab 4.0.1 software (BioRad), respectively. Normalization was done with reference to either GAPDH (housekeeping protein) or respective total protein (EGFR/P38MAPK). Results were reported as mean fold change of target expression in stimulated groups, compared to an unstimulated control group.

### Intracellular imaging of signal pathway activation

HT29 cells were grown in 8-well chambers and treated with LPS and LL-37, either alone or in combination. Cells were fixed, permeabilized using ice-cold acetone (10 min, ^-^20 °C), and rinsed in ice cold PBS-Tw (pH 7.2), followed by blocking in PBS-Tw containing 10% donkey serum, 1% BSA and 0.3 M glycine (1 h, RT). After rinsing with ice cold PBS-Tw, cells were incubated with primary antibodies (16 h, 4 °C) anti-p-p38MAPK (1:3,600). Cells were incubated (1 h, RT) with secondary antibodies Alexa647-conjugated donkey antirabbit IgG (H+L) (711-605-152; Jackson ImmunoResearch) diluted 1:1,000 in 1% BSA in PBS-Tw. Following rinsing in PBS-Tw, cell nuclei were counterstained with DAPI (1:1,000) and mounted with FluorSave. Slides were examined using wide-field immunofluorescence microscope (IX71; Olympus).

### TLR4 and LL-37 knock-down in colonic epithelial cells

A short hairpin (Sh)-TLR4 or (Sh)-LL-37 pGFP-V-RS plasmid vector or noneffective scrambled shRNA construct (sham) (TG320555 and TG314213; Origene) were transfected into confluent HT29 cells using Cell Line Nucleofector^®^ Kit V (VCA-1003; Lonza). Transfected cells were then selected for puromycin resistance (30 μg/mL), followed by fluorescence activated cell sorting (FACS) against green fluorescent protein (GFP). Cell populations (ShTLR4 or ShLL-37-HT29 and sham-HT29) were analysed for TLR4 and LL-37 expression knockdown using western blotting with specific anti-TLR4 antibody (ab22048; Abcam) and anti-cathelicidin antibody (ab87701; Abcam) and LL-37 ELISA (HK321; Hycult Biotechnology), respectively. Knockdown efficiency was ~80-90% and 70-80% for TLR4 and LL-37, respectively, as assessed either through relative intensity quantification of protein bands (for TLR4) or ELISA (for LL-37). Cells were maintained by constantly culturing them in DMEM media supplemented with puromycin (15 μg/mL).

### IL-8 promoter transfection in colonic epithelium and luciferase assay

A pGL3 basic plasmid containing 174bp IL-8 promoter construct (−166 to +8 relative to transcriptional start site) upstream of a firefly luciferase gene and a pRL null plasmid constitutively expressing renilla luciferase gene was utilized (kindly provided by Dr. D. Proud, University of Calgary). The IL-8 promoter construct contained either intact (174 bp full length; FL) or individually mutated sites for NF-κB, activator protein (AP)-1 and nuclear factor for IL-6 expression (NFIL-6) (site-directed mutagenesis; denoted as mNF-κB, mAP-1 and mNFIL-6, respectively). HT29 cells were co-transfected with the promoter construct (1 μg) and pRL (0.1 μg) in DMEM media (FBS and antibiotic free) using TransIT transfection reagent (MIR5405; Mirus Bio LLC) as per manufacturer’s protocol. pRL null plasmid was used as transfection control. After 6 h of transfection, medium was replaced with DMEM containing 10% FBS and 1% penicillin/streptomycin. Cells were then allowed to recover overnight, followed by LPS and LL-37 treatment, in combination or individually. Recombinant human interleukin-1β (IL-1β; 10 ng/mL, 8900; Cell Signaling Technology) was used as a positive control. Results were quantified using a dual luciferase reporter assay (PR-E1910; Promega) as per manufacturer’s instructions. Data were represented as fold change in relative light units (RLU) of firefly luciferase activity with respect to control group, for three independent experiments.

### Human neutrophil isolation and calcium flux assay

Neutrophils were isolated by Lympholyte^®^-poly solution (CL5070; Cedarlane) density gradient centrifugation of heparin-anticoagulated blood, obtained according to ethics-approved procedures from healthy human male donors. Fresh heparin-treated peripheral blood (5 mL) was laid carefully on the top of equal volume of Lympholyte^®^-poly and upon centrifugation (400 x g, 10 min) two bands of leukocytes were apparent. Polymorphonuclear granulocytes were isolated from the lower band and re-suspended in calcium-magnesium free Hank’s Balanced Salt Solution (HBSS; Gibco, Life Technologies). For calcium flux assay, neutrophils were incubated with Fluo4-no wash calcium dye (Fluo4 NW, F1242; Invitrogen) (1 mL; 45 min, RT), washed with PBS (1x), and re-suspended in calcium-magnesium containing HBSS. Neutrophils (1 x 10^6^) were transferred into a cuvette and stimulated with supernatants (100 μL) harvested from HT29 cells, either untreated (control; serum free DMEM media) or treated with LPS ± LL-37. To confirm the role of IL-8, supernatants from HT29 cells treated with LPS ± LL-37 were blocked with anti-IL-8 antibody (1 μg/mL, MAB208; R&D Systems), CXC receptor1/2 (CXCR1/2) inhibitor (SCH 527123, 20 μM, A3802; APExBIO), or mouse IgG1 isotype control (1 μg/mL, 5415S; Cell Signaling Technology) (1 h, 4 °C) prior to stimulation of neutrophils. Recombinant human IL-8 (200 ng/mL, 208-IL-010; R&D Systems) was used as positive control. Calcium signals were monitored at an excitation wavelength of 480 nm and an emission wavelength of 530 nm, recorded with the Aminco Bowan series II fluorimeter and AB2 software (Thermo Fisher Scientific). Data were expressed as a percentage of the emission fluorescence normalized to maximum fluorescence caused by calcium ionophore (CI A23187, 2.5 μM, 52665-69-7; Sigma) per experiment, for a total of three independent experiments.

### Neutrophil activation by assessment of neutrophil elastase secretion

To assess the role of IL-8 secreted by colonic epithelial cells in the activation of neutrophils, human neutrophils were isolated as discussed above, seeded in 8-well chambers (1 x 10^4^ cells/well) and stimulated with supernatants (500 μL) harvested from HT29 cells, either untreated (control; serum free DMEM media) or treated with LPS ± LL-37 (for 4 h). Neutrophils were fixed in 4% PFA (15 min, RT) and rinsed in ice cold PBS-Tw (pH 7.2), followed by blocking in PBS-Tw containing 10% donkey serum, 1% BSA and 0.3 M glycine (1 h, RT). After rinsing with ice cold PBS-Tw, neutrophils were incubated with primary antineutrophil elastase antibody (5 μg/mL; MAB91671; R&D Systems) diluted in PBS (16 h, 4°C) and then, incubated with secondary Alexa549-conjugated donkey anti-mouse IgG (H+L) antibodies diluted (1: 1,000) in 1% BSA in PBS-Tw (1 h, RT). Neutrophils were rinsed with PBS-Tw, counterstained with DAPI (1:1,000), and mounted with FluorSave. Slides were examined using a wide-field immunofluorescence microscope (IX71; Olympus). For neutrophil elastase secretion quantification, fluorescence intensity was calculated using ImageJ 1.50i software. Average fluorescence intensity per replicate was quantified in five randomly selected fields of view (~5-6 neutrophils/field of view). Each experiment was done in triplicate and MFI for an experiment was obtained by averaging total of 15 fields of view (five per replicate; total of 75-90 neutrophils per experiment) and represented as a single numerical value. Data were MFI normalized to their respective control for three independent experiments and represented as fold increase in MFI.

### LPS uptake assay

HT29 cells grown in 8-well chamber were treated with Alexa 488-conjugated-LPS ± LL-37 peptides. Cells were fixed with 4% PFA in the dark (15 min, RT), rinsed in ice-cold PBS-Tw (3 times, pH 7.2), counterstained for nuclei with DAPI (1:1,000) (30 min, RT), rinsed with PBS-Tw (3 times), and mounted with FluorSave reagent. Slides were examined using wide-field immunofluorescence microscope (IX71Olympus). Assessment of Alexa 488-conjugated-LPS uptake by colonic cells was performed using ImageJ 1.50i software. Fluorescence intensity was calculated either in randomly selected fields of view with a cluster of ~8-10 cells (for time-dependent LPS uptake assay,) or ~8-10 single cells (for LPS uptake in presence of inhibitors) per replicate, for a total of three replicates/experiment. A single numerical value (average of three replicates) was obtained for individual experiments and reported as MFI for three independent experiments.

### Location of TLR4 in colonic epithelial cells

HT29 cell monolayers were washed with warm PBS (1x) and lifted with 0.25% trypsin-EDTA. Cells were centrifuged (300 x g, 5 min), re-suspended in stain buffer (FBS, 554656; BD Bioscience), and fixed either with BD Cytofix (554655; BD Bioscience) for surface staining or BD Cytofix/Cytoperm (554717; BD Bioscience) for intracellular staining (20 min, room temperature). Cells were blocked with human BD Fc Block (564219; BD Bioscience) (2.5 μg/mL, 10 min, RT), washed as described above, and incubated with either PE labelled anti-TLR4-CD284 (12-9917-41; eBioscience) or PE labelled isotype mouse IgG (12-4724-41; eBioscience) (0.5 μg/mL, 30 min, 4°C). Cells were then suspended in flow buffer (0.4 mL), passed through fluorescence-activated cell sorting (FACS) machine (BD^TM^ LSR II; BD-Bioscience) and gated for the single cell population that was PE positive. Data were obtained as mean fluorescence intensity (MFI) and represented as ratio of intracellular to surface TLR4 expression.

### Ethics statement

All animal studies were reviewed and approved by the University of Calgary Animal Care Committee that adheres to the principles and policies in the “Guide to the Care and Use of Experimental Animals” published by the Canadian Council on Animal Care (Animal Protocol # AC16-0092). For human colonoids, five colonic biopsies per patient were obtained upon written consent from two informed, unidentified healthy patients undergoing routine prophylactic cancer surveillance colonoscopy. All procedures were done in accordance with protocols approved by the University of Calgary Cumming School of Medicine Human Ethics Committee (REB18-0104).

### Statistical analyses

Analytical data represented as histograms were recorded as mean values with bars representing standard errors of the mean (SEM) from a minimum of three independent experiments, with data obtained in triplicates, unless otherwise mentioned. Normality was assessed using D’Agostino & Pearson omnibus normality or Shapiro-Wilk (Royston) tests. All comparisons were performed using either two-sided unpaired Student’s *t*-test or one-way analysis of variance (ANOVA) with a *post hoc* Bonferroni correction for multiple group comparisons. A *P*-value was assigned to each group with reference to control group, unless shown specifically on the graph. A *P* value of <0.05 was considered significant. All statistical analysis was performed with Graph Pad Prism software (Graph Pad 5.0).

## Results

### Cathelicidin LL-37 acts synergistically with S. typhimurium via TLR4 to stimulate IL-8 synthesis in colonic epithelium

LL-37, the only endogenous cathelicidin in humans, induces IL-8 synthesis in airway and corneal epithelial cells (30, 31). Furthermore, *S. typhimurium* promotes IL-8 secretion in colonic epithelial cells (HT29 and T84) via NF-κB-dependent pathways (32). Whether LL-37 can synergize with *S. typhimurium* to modulate IL-8 cytokine expression by colonic epithelia was uncertain. Thus, we first studied effects of exogenous cathelicidin, to determine if HT29 cells challenged by *S. typhimurium* and stimulated with synthetic LL-37 peptide had enhanced production of IL-8 (Fig 1A). There were time-dependent increases in colonic IL-8 production (up to 4 h, with no significant difference at 8 h) (Fig 1A (i)) under various LL-37 concentrations (1 to 20 μg/mL) (Fig 1A (ii)). LL-37 alone at concentrations up to 30 μg/mL did not induce IL-8 production (data shown for 4 h) (S2A Fig). To examine mechanisms of exogenous cathelicidin to stimulate IL-8 in synergy with *S. typhimurium,* LL-37 at 10 μg/mL for 4 h was used as our experimental treatment. shRNA-mediated knock-down of TLR4 (Fig 1B (i)), established Gram-negative bacterial receptor, diminished the ability of *S. typhimurium,* acting either on its own or along with LL-37 to enhance IL-8 secretion (Fig 1B (ii)). Next, we investigated the role of naturally occurring cathelicidin secreted by colonic epithelium in synthesis of IL-8, as secretion of endogenous cathelicidin may be an epithelial defense mechanism. *S. typhimurium* induced synthesis and peptide secretion of LL-37 in HT29 cells over a 2-8 h interval (Figs 1C (i-ii)). Furthermore, endogenous cathelicidin in the colonic epithelium promoted IL-8 synergistically with *S. typhimurium,* as secretion of IL-8 diminished in gene-specific knockdown of LL-37 in HT29 cells stimulated by *S. typhimurium* (Fig 1C (iii)). When identifying *S. typhimurium* bacterial factor(s) involved in this synergistic IL-8 induction, combined stimuli with LL-37 and LPS was found to enhance secretion of IL-8 in HT29 cells (Fig 1D). Intriguingly, the lipid A component of LPS reduced the constitutive production of IL-8 by HT29 cells and did not augment IL-8 when combined with LL-37 (Fig 1E).

**Fig 1.**
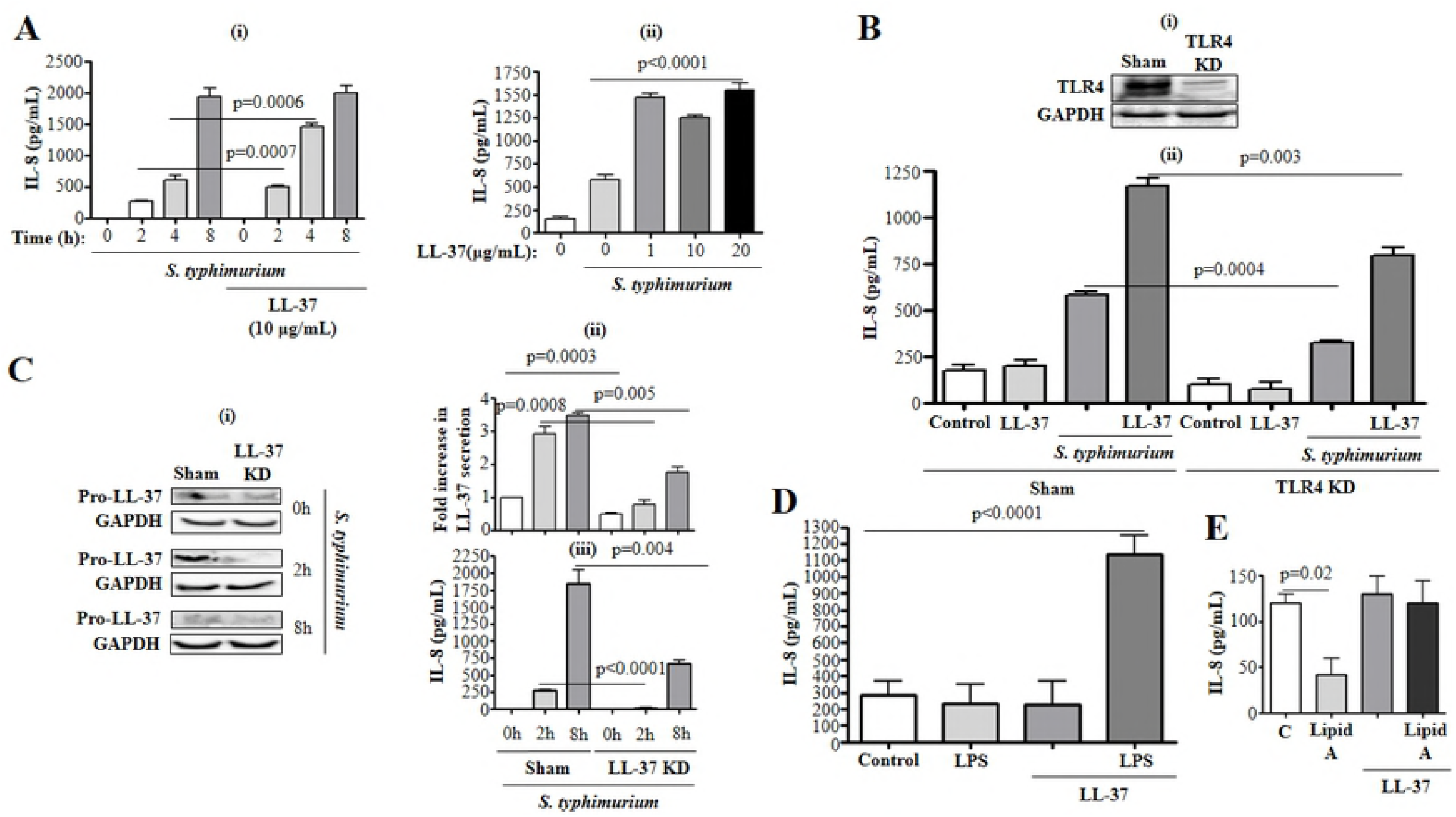
TLR4 dependent IL-8 synthesis in colonic epithelial cells upon combined stimulation with cathelicidin and either *S. typhimurium* or derived LPS. **(A-C)** HT29 cells either **(A)** normal or **(B-C)** sham transfected/knocked down for **(B *i*)** TLR4 or **(C)** LL-37, were either stimulated with 10 μg/mL (**A *i* and B *ii***) or variable concentrations (1 to 20 μg/mL) (A ii) of synthetic LL-37 or not (C), followed by challenge with *S. typhimurium* (MoI 10:1; 2 x 10^7^ CFU/mL) for variable time points (**A *i and C***) or 4 h (**A *ii* and B *ii***), and subsequent quantification of secreted IL-8 using ELISA (**A, B *ii* and C *iii***). (**C *i-ii***) LL-37 protein expression in cell lysate (**C *i***) and cell supernatant (C *ii*) was determined using western blotting and ELISA, respectively. (**A-C**) Figures represent one of three independent experiments. (**D-E**) HT29 cells were challenged with (**D**) LPS (1 μg/mL) or (**E**) lipid A (1 μg/mL) and LL-37 (10 μg/mL), either alone or in combination, for 4 h, followed by quantification of IL-8 in cell supernatants using ELISA. Data are shown as means ± SEM (*n* = 3 independent experiments done in triplicate). A *p*-value of < 0.05 (one-way ANOVA, with *post hoc* Bonferroni correction for multiple group comparison or two-tailed Student’s *t*-test for two groups) were considered significant.

To understand the biological relevance of IL-8 secreted by the colonic epithelium when sensing enteric pathogens in the presence of cathelicidin, secreted products released by HT29 cells in response to LPS/LL-37 were exposed to naïve human neutrophils. Supernatants from colonic epithelial cells exposed to LPS/LL-37 activated human neutrophils, as assessed by calcium flux (S3A Fig), and promoted secretion of neutrophil elastase (S3B Fig). This response was mostly attributed to IL-8, as anti-IL-8 antibody and an IL-8 receptor antagonist blocked both events (S3A and S3B Figs). Therefore, LL-37 and either intact *S. typhimurium* or LPS (but not lipid A) synergistically stimulated production of IL-8 by colonic epithelial cells. IL-8 production was partially dependent on the presence of TLR4 (Fig 1B). In addition, IL-8 secreted by colonic epithelium when stimulated by cathelicidin and *S. typhimurium* seemed to promote neutrophil activation.

### Coordinated synergy of exogenous cathelicidin and LPS promotes IL-8 in the colonic epithelium of humans and mice

Based on studies of IL-8 kinetics, upregulation of *IL-8* mRNA was optimal at 2 h but declined thereafter to an apparent ‘steady-state’ (significantly above baseline) at 3-4 h after LL-37/LPS treatment (Fig 2A). Secretion of IL-8 began as early as 1 h after LL-37/LPS treatment and continued rising until 16 h post-stimuli (Fig 2B). Based on 4 h as a time point for measuring IL-8 secretion, concentration-dependence of the synergistic actions of LL-37 and LPS was evaluated using: 1) increasing concentrations of LPS for a fixed LL-37 concentration (10 μg/ml; ~ 2 μM) (Fig 2C(i)); and 2) increasing concentrations of LL-37 for a fixed amount of LPS (1 μg/mL) (Fig 2C(ii)). A bell-shaped biphasic concentration-response curve was observed for combined LL-37 and LPS, with optimal concentrations of 1 μg/mL for LPS (Fig 2C(i)) and 3 μg/mL for LL-37 (Fig 2C(ii)). Neither LPS nor LL-37 alone had concentration-related increases in IL-8 production as demonstrated by their combinatorial action (S2B and S2A Figs). To assess whether LL-37/LPS modulates IL-8 at transcription or translational levels, actinomycin-D was used to block transcription and cycloheximide to block protein translation. Both actinomycin-D and cycloheximide inhibited secretion of IL-8 in response to LPS/LL37 (Fig 2D). Thus, LL-37/LPS seemed to promote *a novo* synthesis (transcription and translation) of IL-8 in colonic epithelium.

**Fig 2.**
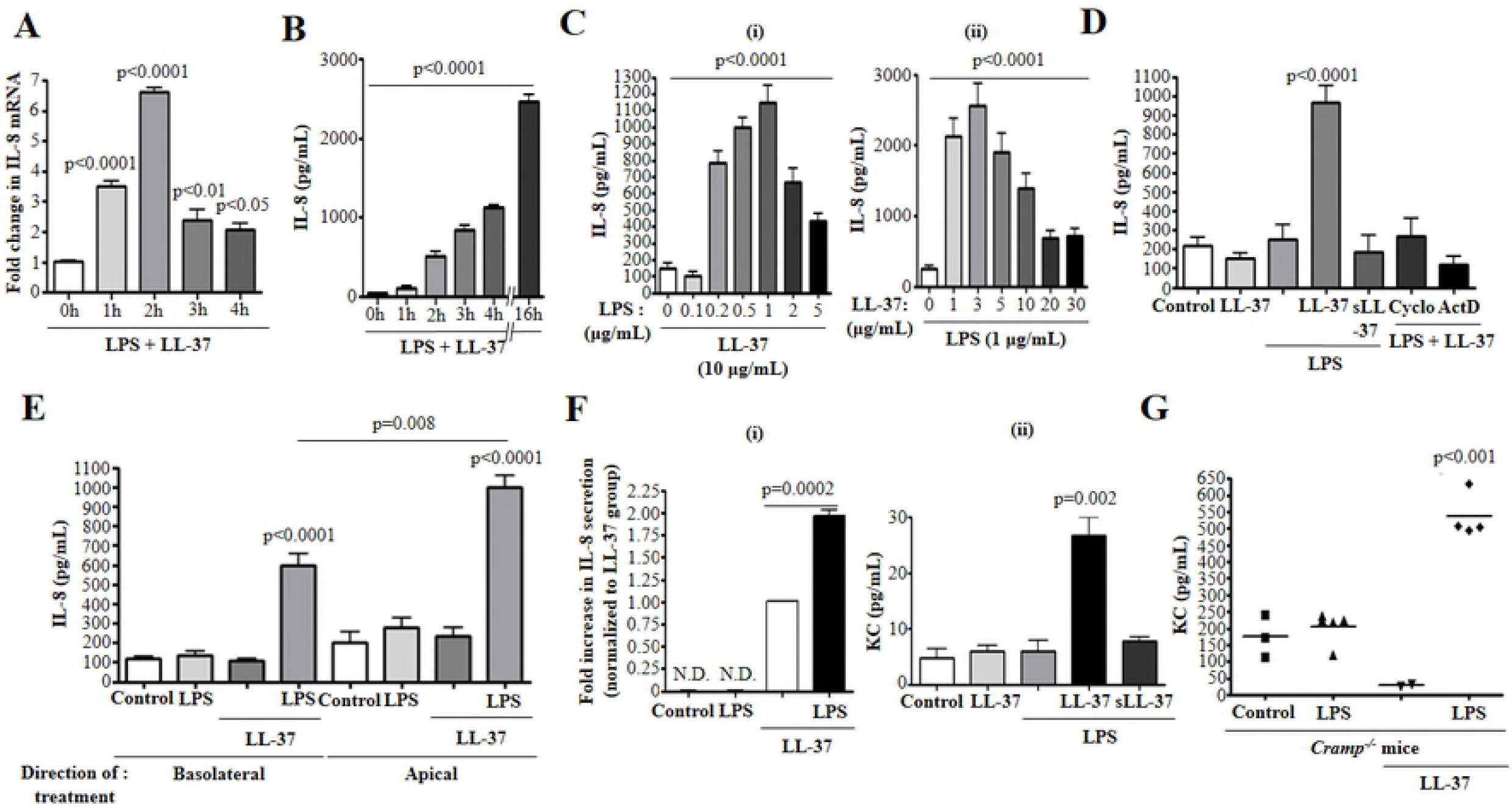
Synthetic cathelicidin and LPS induced IL-8 synthesis in colonic epithelia, mouse colonoids, and cathelicidin deficient mice. **(A-B)** HT29 cells were stimulated with LPS (1 μg/mL) and LL-37 (10 μg/mL) for various intervals. (**C *i-ii***) HT29 cells were challenged with either variable dose of LPS and constant LL-37 (10 μg/mL) (**C *i***), or variable dose of LL-37 and constant LPS (1 μg/mL) (n= 4) (**C *ii***), followed by IL-8 quantification at **(A)** mRNA (n=4) and **(B-C)** protein levels using qPCR and ELISA, respectively. **(D)** HT29 cells were pre-treated or not, with either a protein synthesis inhibitor, cycloheximide (Cyclo; 1 μM), or a transcriptional inhibitor, actinomycin D (ActD; 1 μM) for 1 h, followed by stimulation with LPS, LL-37, or scrambled peptide (sLL-37; 10 μg/mL), either alone or in combination, for 4 h. IL-8 protein secretion was quantified in cell supernatant by ELISA. **(E-G) (E)** T84 cells, **(F)** colonoids from (**F *i***) human patients (five biopsies/patient, from two patients), or (**F *ii***) from wild type *(Cramp^+/+^)* mice and (G) *Cramp^1-^* mice (*n*= 2-5 mice/per group) were treated (*ip* for **(G)**) with LPS, LL-37, or scrambled-sequence peptide, sLL-37 (only in mouse colonoids) either alone or in combination, for 4 h (3 h for *Cramp^−/−^* mice). IL-8 protein secretion in supernatant (**F *i***), apical compartments (**(E)** and (**F *ii***)), or mouse colonic tissue **(G)** was quantified by ELISA. (**F *i***) Data were normalized to LL-37 treatment group as IL-8 secretion in control and LPS-only treatments were below detection. Data were represented as fold increase in IL-8 secretion to minimize inter-patient variability. Data are shown as mean ± SEM (*n* = 3 independent experiments done in triplicate, unless mentioned otherwise). *P* < 0.05 (one-way ANOVA*post hoc* Bonferroni correction for multiple group comparison or two-tailed Student’s *t*-test for two groups) was considered significant. ND = ‘not detected’.

The synergistic action of LL-37 with LPS was further investigated in another colonic epithelial model, colonoid systems, and *in vivo* murine models. These studies were conducted using LL-37 at 10 μg/mL and LPS at 1 μg/mL, based on previous experiments (Figs 2C(i-ii)). Apical-basal polarized colonic epithelial T84 cells exposed to LPS/LL-37 secreted IL-8 into the apical compartment, irrespective of the direction of stimuli (Fig 2E). This cooperation between LL-37 and LPS was extended to *in vitro* human and mouse-derived colonoids cultured for 10 d when they displayed a morphology compatible with large intestinal crypts (S1 Fig). A combination of LL-37/LPS (but not individually) stimulated cytokine secretion (IL-8 for human tissues; KC for murine tissues) in both human (Fig 2F (i)) and polarized mouse colonoids (Fig 2F (ii)) for which apical secretion of KC was detected. Treatment with scrambled sequence peptide (sLL-37) did not synergize with LPS to augment IL-8 in HT29 cells (Fig 2D) or KC secretion by mouse colonoids (Fig 2F (ii)). To further validate the ability of LL-37 to synergize with LPS at colonic mucosa, cathelicidin-deficient Cramp-null mice *(Cramp^−/−^)* (13) were given LPS ip and supplemented with synthetic LL-37 to reconstitute a cathelicidin effect (Fig 2G). Systemic injection of combined LL-37 and LPS to *Cramp^−/−^* mice increased colonic KC at 3 h (Fig 2G). LPS and LL-37 alone did not significantly change tissue content of KC (Fig 2G). Thus, cathelicidin and LPS synergistically triggered production of chemoattractant cytokine (IL-8/KC) in intestinal-derived cells and colons.

### LL-37/LPS interaction-mediated and TLR4 dependent-enhanced IL-8 synthesis in colonic epithelium

Binding of LPS to its natural receptor TLR4 increases IL-8 synthesis in intestinal cells (33) and TLR4 over-expressing kidney epithelial cells (HEK293) (34). In this study, induction of IL-8 by the combined cathelicidin and *S. typhimurium* was in part dependent on colonic epithelial TLR4, as observed in shRNA-mediated TLR4 knockdown HT29 cells (Fig 1C (ii)). This necessity of colonic epithelial TLR4 was also confirmed for the combined action of LL-37 and LPS in increasing IL-8. The TLR4 antagonist, LPS-RS, inhibited upregulation of *IL-8* mRNA and protein in HT29 cells upon stimulation with LL-37/LPS (Figs 3A (i) and (ii)). Likewise, shRNA-mediated knockdown of TLR4 in HT29 cells reduced IL-8 secretion in response to LPS/LL-37 (Fig 3B).

**Fig 3.**
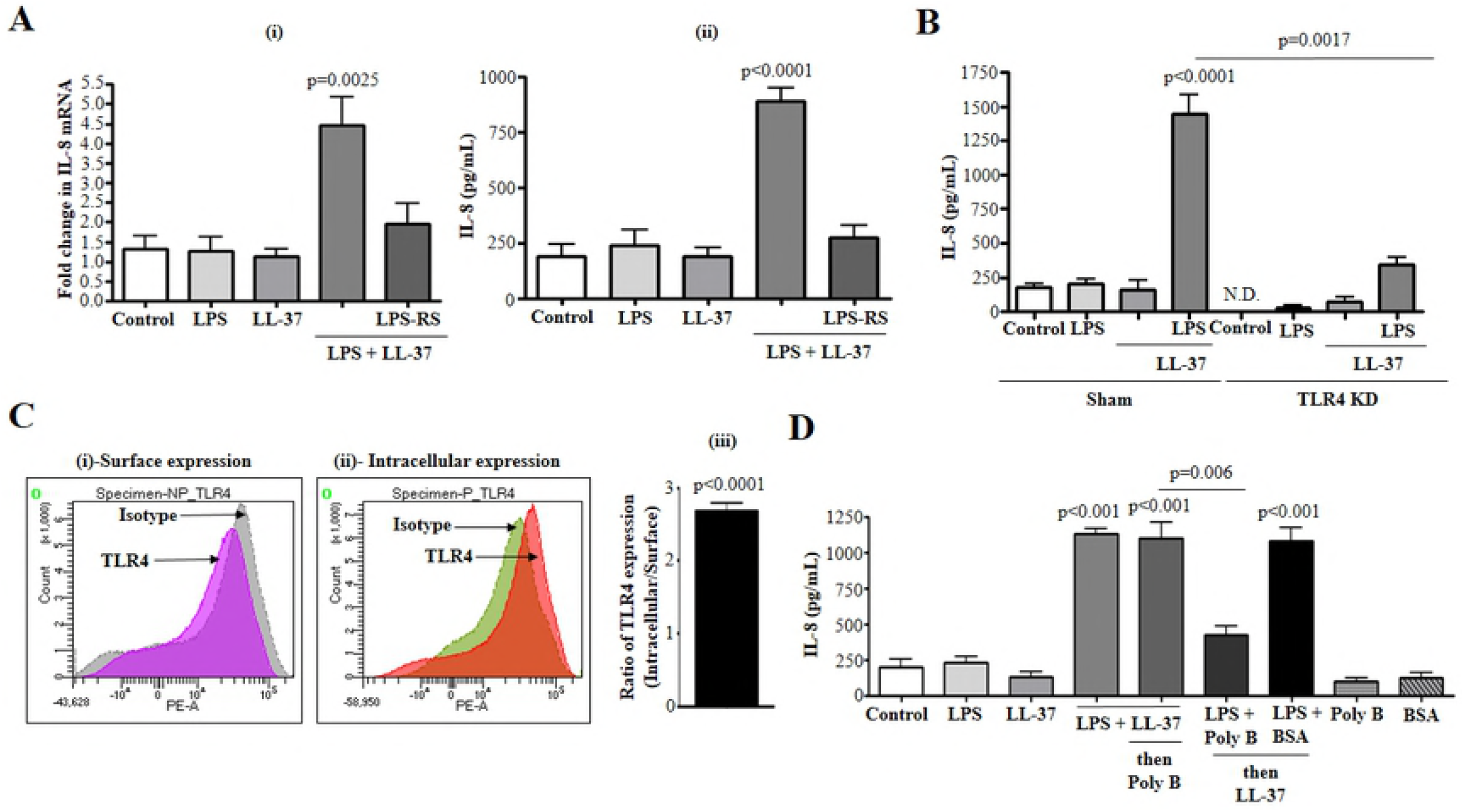
LPS-LL-37 interaction and intracellular TLR4 dependent IL-8 synthesis in colonic epithelial cells. (**A *i-ii***) HT29 cells were pre-treated or not with LPS-RS (5 μg/mL) for 1 h, followed by stimulation with LPS and LL-37 either alone or in combination, for 1 h **(A *i*)** and 4 h **(A *ii*)**. **(A *i*)** *IL-8* mRNA and **(A *ii*)** protein secretion (n= 4) were quantified using qPCR and ELISA, respectively. **(B)** HT29 cells either sham transfected or knocked down for TLR4 (TLR4 KD), were challenged with LPS (1 μg/mL) and LL-37 (10 μg/mL) either alone or in combination, for 4 h. **(C)** Flow cytometric assessment of TLR4 expression in non-permeabilized (surface expression) (**C *i***) or permeabilized (intracellular expression) (**C *ii***) HT29 cells using PE-labelled anti-TLR4-CD284 antibody (0.5 μg/mL). Images are representative of three independent experiments. Isotype mouse IgG was used as a negative control. (**C *iii***) Histogram representing ratio of intracellular to surface TLR4 expression in HT29 cells per experiment. **(D)** HT29 cells were treated with LPS and LL-37, either alone or in combination, or pre-incubated with a mix of LPS (1 μg/mL)/LL-37 (10 μg/mL) (1 h, 37°C) followed by substitution with polymyxin B (Poly B; 10 μg/mL), or pre-incubated with a mix of LPS (1 μg/mL)-polymyxin B (Poly B; 10 μg/mL)/bovine serum albumin (BSA;10 μg/mL) (1 h, 37°C) and substituted with LL-37 (10 μg/mL). IL-8 secretion was quantified after 4 h using ELISA. Polymyxin B alone and BSA were used as negative controls. Data are shown as means ± SEM (n = 3 independent experiments done in triplicate). *P* < 0.05 (one-way ANOVA *post hoc* Bonferroni correction for multiple group comparison or two-tailed Student’s *t*-test for two groups) was considered significant. ND = ‘not detected’.

Since TLR4 was critical for LL-37/LPS-induced IL-8 prodution, we next sought to determine the cellular location of TLR4 in colonic epithelium. Based on flow cytometery, HT29 cells had a constitutively low level of TLR4, mainly intracellular with negligible cell surface expression (Figs 3C (i-iii)). Therefore, we questioned how cathelicidin interacted with LPS and gained access to internal TLR4. Cathelicidins are cationic peptides that bind to negatively charged molecules such as LPS. LPS pre-treated with polymyxin B (a cyclic cationic polypeptide antibiotic that binds to and neutralizes LPS) was not able to synergize with LL-37 for IL-8 synthesis in HT29 cells (Fig 3D). However, such an inhibitory effect was not observed with addition of bovine serum albumin (BSA; negative control) (Fig 3D). In addition, synergistic IL-8 production did occurr if LPS was incubated with LL-37 prior to addition of polymyxin B (Fig 3D). Thus, a pre-formed extracellular LPS-LL37 complex seemed to gain access to intracellular TLR4 as a first step in IL-8 synthesis in colonic epithelium.

### IL-8 synthesis stimulated by LPS and cathelicidin in colonic epithelium depends on monosialotetrahexosylganglioside (GM1)-lipid raft mediated LPS uptake

Based on bioinformatic analysis, LL-37 shared structural homology with the cholera toxin B-subunit B (CtxB: S4 Fig). CtxB internalizes the active toxin A-subunit by binding to ganglioside GM1 (monosialotetrahexosylganglioside) (35), a main lipid raft component in the plasma membrane (36). We therefore hypothesized that GM1-containing lipid rafts aid in internalizing the LL-37/LPS complex and interacting with intracellular TLR4. As expected, there was a time-dependent increase in Alexa 488-conjugated-LPS uptake in presence of LL-37 in HT29 cells (Fig 4A). When lipid rafts were disrupted by methyl-β-cyclodextran (MBC; 500 μM) or by blocking cholesterol synthesis with mevalonate (Mev: 250 ng/mL), the ability of LPS/LL-37 to induce IL-8 secretion (Fig 4B) and internalization of Alexa 488-conjugated-LPS (Fig 4C) were abolished. Similarly, addition of exogenous GM1 (100 μg/mL) to compete with LL-37/LPS complexes in the access to lipid rafts blocked IL-8 secretion in HT29 cells (Fig 4D). Other endocytic pathways, including dynamin GTPase-dependent or actin-dependent processes assessed by pharmacological inhibiton using dynamin inhibitor D15 or cytochalasin D, were not involved in uptake of Alexa 488-conjugated-LPS/LL-37 complex by HT29 cells (Figs 4B, C). Thus, LPS-LL-37 synergism in promoting IL-8 production by colonic epithelial was dependent on extracellular binding with GM1 and lipid raft-mediated uptake.

**Fig 4.**
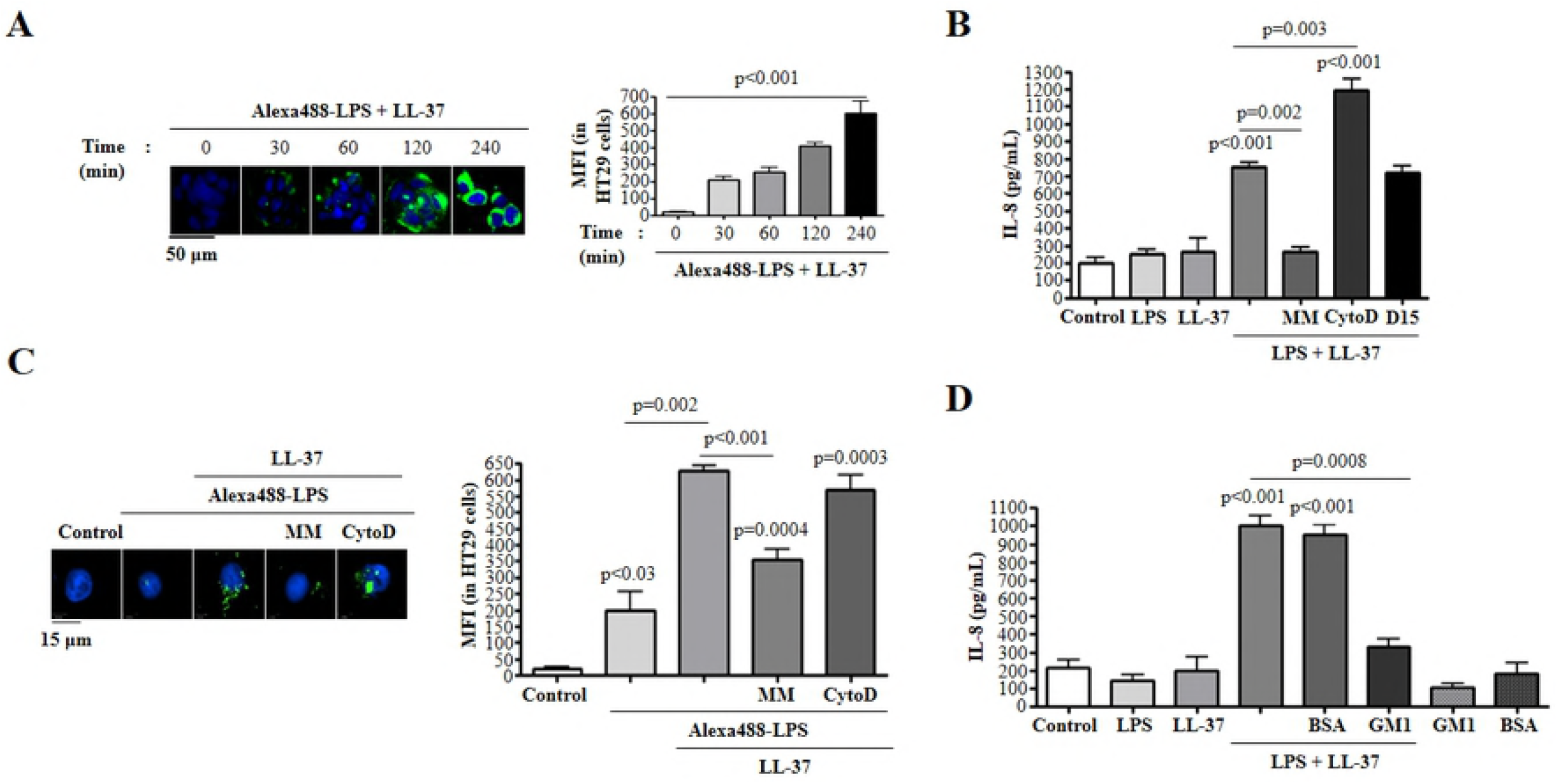
LL-37 mediated LPS uptake regulate IL-8 synthesis in colonic epithelial cells. **(A)** HT29 cells were challenged with a combination of Alexa 488-conjugated-LPS (1 μg/mL) and LL-37 (10 μg/mL) for variable interval (0, 30, 60, 120 and 240 min), followed by quantification of fluorescence using ImageJ 1.50i software. Fluorescence was calculated and represented as mean fluorescence intensity (MFI) for three independent experiments. **(B, C)** HT29 cells were pre-treated with a mix of lipid raft inhibitors (named MM, cocktail of methyl-β-cyclodextrin (500 μM) and mevinolin (250 ng/mL)) or endocytosis inhibitors cytochalasin D (CytoD; 2 μM) or dynamin-dependent endocytosis inhibitor (D15; 10 μM) for 1 h, followed by stimulation with **(B)** LPS (1 μg/mL) or **(C)** Alexa 488-conjugated-LPS (1 μg/mL) and LL-37 (10 μg/mL), either alone or in combination, for 4 h. **(B)** IL-8 protein secretion was quantified using ELISA. **(C)** LPS uptake was assessed using immunocytochemistry and quantified using ImageJ 1.50i software. Fluorescence was calculated and represented as mean fluorescence intensity (MFI) for three independent experiments. **(D)** IL-8 was quantified by ELISA in supernatants from HT29 cells treated with LPS and LL-37 in presence (or not) of exogenous monosialotetrahexosylganglioside (GM1; 100 μg/mL). BSA (100 μg/mL) was used as a negative control. Data are shown as means ± SEM (n = 3 independent experiments, done in triplicate). *P* < 0.05 (one-way ANOVA*post hoc* Bonferroni correction for multiple group comparisons or two-tailed Student’s *t* test for two groups) were considered statistically significant.

### Cathelicidin mediated IL-8 synthesis by colonic epithelium in response to LPS requires EGFR kinase activity

Intracellular signalling for IL-8 biosynthesis in colonic epithelium involves protein tyrosine kinases, MAPKs, and NF-κB (37, 38) although their detailed interactive pathway is not fully understood. Studies using human airway epithelial cells demonstrated a key role of EGFR transactivation in IL-8 production induced by LL-37, along with a TLR1/2 agonist (Pam3CSK4) (39, 40). In our study, phosphorylation of EGFR increased after LPS/LL-37 over 2 h (Fig 5A); this was stronger than effects with a single stimuli (i.e., control, LPS or LL-37) (data shown for 2 h; Fig 5B). EGFR kinase activity was not required for IL-8 gene transcription but determinant in protein synthesis as inhibition of EGFR kinase activity (AG1478) did not affect LPS-LL-37 induced *IL-8* mRNA levels (Fig 5C) but blocked IL-8 secretion (Fig 5D).

**Fig 5.**
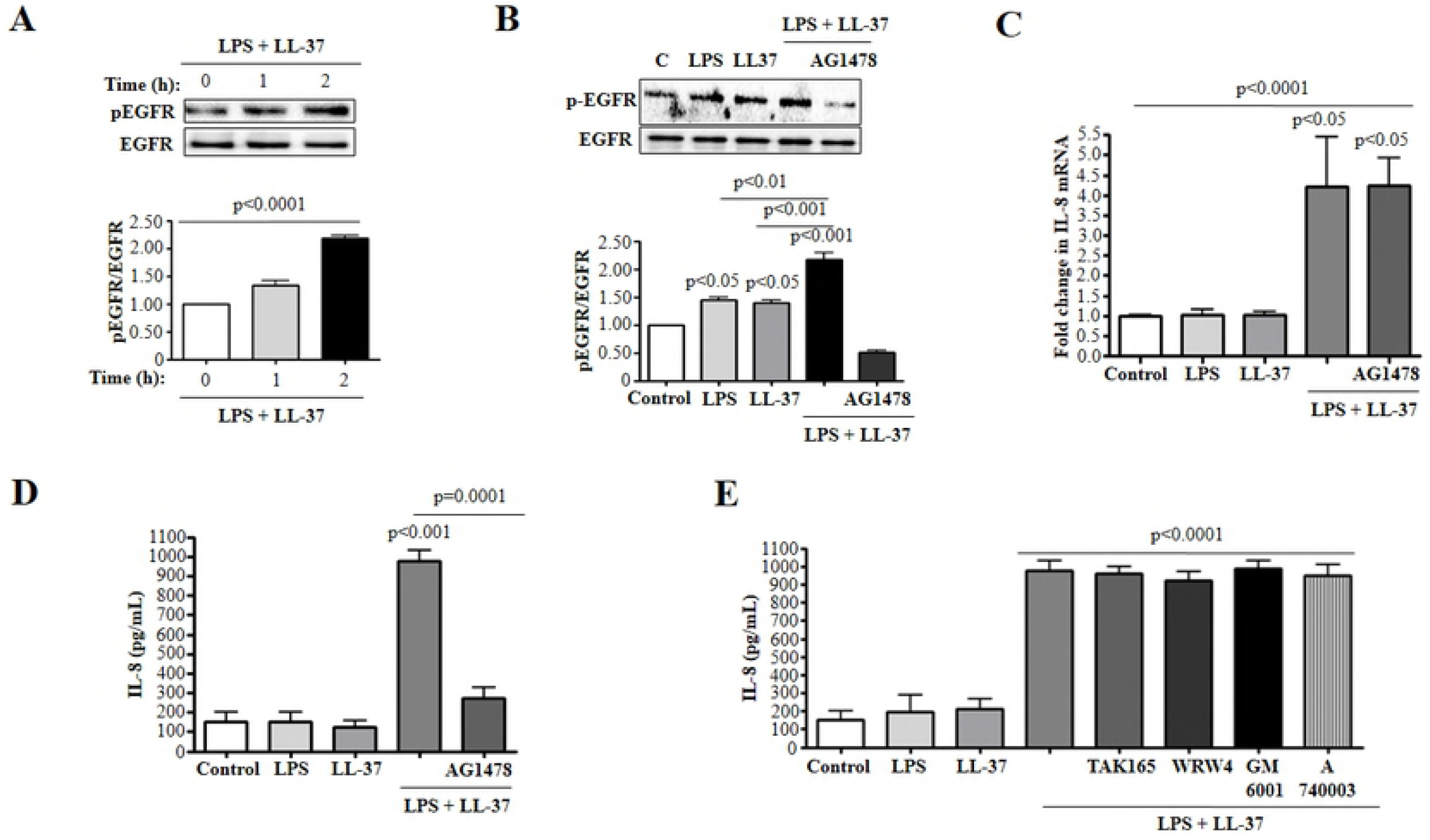
EGFR signalling promotes IL-8 secretion in colonic epithelial cells stimulated by cathelicidins and LPS. **(A-D)** HT29 cells were pre-treated **(B-D)** or not **(A)** with AG1478 (1 μM) for 1 h, followed by treatment with LPS (1 μg/mL) and LL-37 (10 μg/mL), either **(B-D)** alone or in **(A-D)** combination. **(A)** Time dependent or **(B)** at 2 h activation of EGFR in colonic cells (HT29) was assessed by western blotting with antibodies against phosphorylated Tyr-1068 EGFR. Total loading was confirmed after immunoblotting with EGFR. **(C)** *IL-8* mRNA synthesis was quantified after 1 h using qPCR (n=4) and **(D)** protein secretion (n=4) was assessed after 4 h using ELISA. **(E)** HT29 cells were pre-treated with inhibitors TAK165 (ErbB2; 10 μM), WRW4 (FPRL1; 10 μM), GM6001 (MMP; 10 μM), and A740003 (P2X7; 10 μM) or not, followed by treatment with LPS and LL-37, either alone or in combination for 4 h (n= 4). IL-8 secretion was quantified using ELISA. Data are shown as means ± SEM (n = 3 independent experiments done in triplicate). *P* < 0.05 (one-way ANOVA *post hoc* Bonferroni correction for multiple group comparison or two-tailed Student’s *t*-test for two groups) were considered statistically significant.

LL-37 signals through distinct membrane based receptors, including FPRL-1 (41), P2X7 (42), and ErbB2 (43) in neutrophils, macrophages and skin-derived epithelial cells. However, we found that pharmacological inhibition of FPRL-1, ErbB2, and P2X7 receptors did not affect synergistic actions of cathelicidin and LPS to stimulate IL-8 secretion (Fig 5E). Therefore, production of IL-8 stimulated by LPS/LL-37 relied on post-transcriptional effects of EGFR kinases, whereas it was independent of FPRL-1, ErbB2 and P2X7 receptors.

### IL-8 synthesis stimulated by LPS/LL-37 in colonic epithelium signals through TLR4/Src kinase and p38MAPK activation

Immunomodulatory activities of LL-37 in a variety of epithelia, including those isolated from the eye (31) and airway (30), is dependent on MAPK signalling (both ERK and p38MAPK). We detected time-dependent activation of p38MAPK by LPS/LL-37 in HT29 cells (Figs 6A and S5 Fig), whereas inhibition of p38MAPK (SB203580) reduced the LPS/LL-37 mediated upregulation of *IL-8* mRNA and protein (Figs 6B (i) and (ii)). However, inhibition of ERK1/2 (FR180204) did not affect IL-8 secretion (Fig 6B (ii)). It has been reported that EGFR phosphorylation mediates the p38MAPK activation of the cyclooxygenase 2 (COX-2) and vascular endothelial growth factor (VEGF) expression in human pancreatic duct epithelial cells (44). In agreement, we showed EGFR-kinase inhibition (AG1478) reduced phosphorylation of p38MAPK (Fig 6C) and blocked IL-8 secretion induced by LPS and LL-37 (Figs 5D and 6B (ii)).

**Fig 6.**
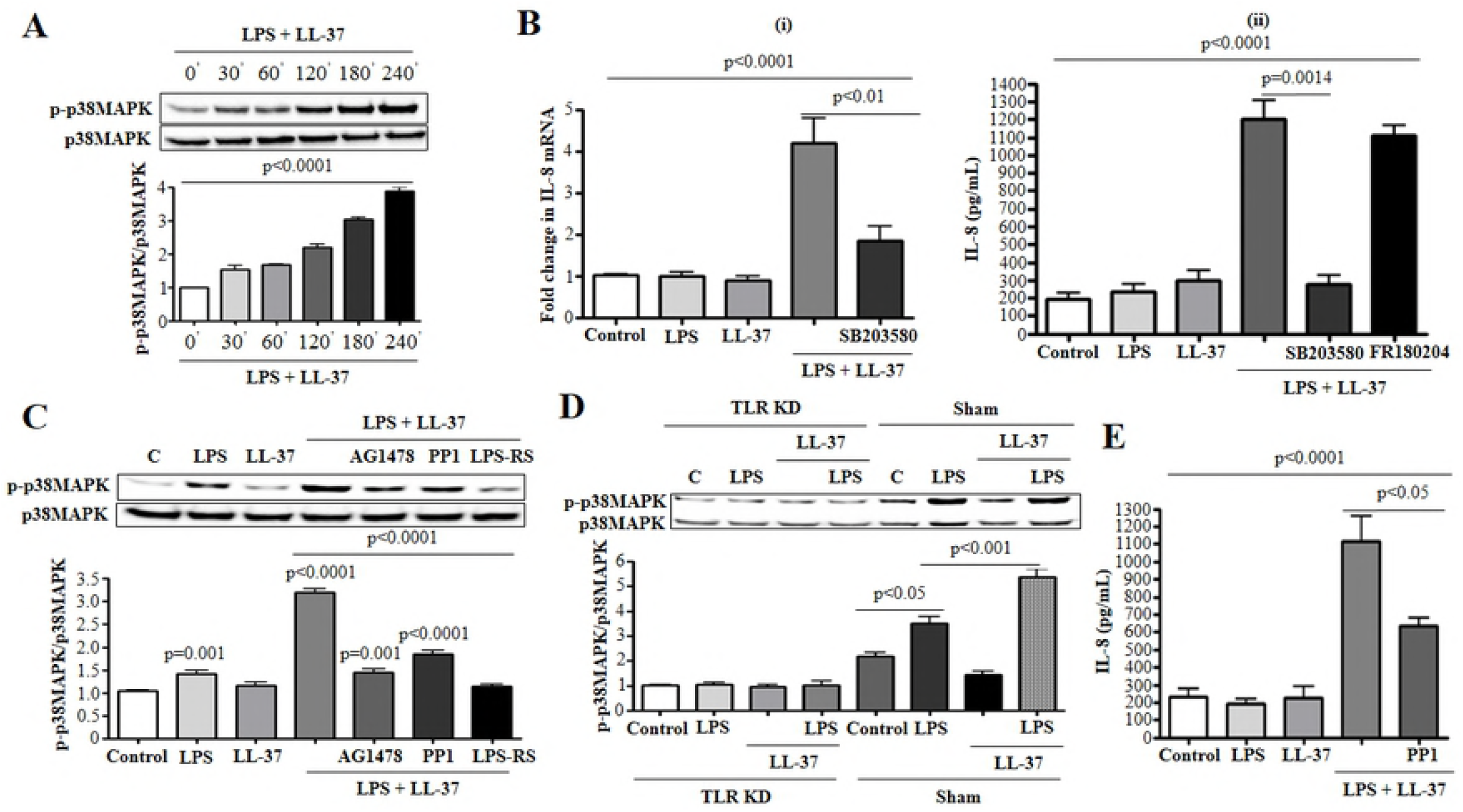
p38MAPK promotes IL-8 synthesis in colonic epithelial cells stimulated by cathelicidin and LPS. **(A)** LPS-LL-37 mediated, time-dependent activation of p38MAPK in HT29 cells was confirmed using western blotting with antibodies against phosphorylated p38MAPK. (B) HT29 cells were pre-treated with (**B *i-ii***) p38MAPK inhibitor SB203580 (2 μM) or (**B *ii***) ERK inhibitor FR180204 (10 μM) for 1 h, followed by treatment with LPS (1 μg/mL) and LL-37 (10 μg/mL), either alone or in combination. **(B *i*)** *IL-8* mRNA synthesis was quantified after 1 h using qPCR (n=4) and, (**B *ii***) protein secretion was assessed after 4 h using ELISA. **(C)** HT29 cells treated with inhibitors for EGFR kinase, Src kinases or TLR4, denominated AG1478 (1 μM), PP1 (1 μM) or LPS-RS (5 μg/mL), respectively, for 1 h, or **(D)** HT29 cells either sham transfected or knocked down for TLR4 (TLR4 KD), followed by LPS and LL-37 treatment, either alone or in combination for 2 h. p38MAPK phosphorylation was assessed by western blotting with specific antibodies. Total loading was confirmed after blotting for p38MAPK. (E) HT29 cells were pre-treated with Src kinases inhibitor PP1 (1 μM) followed by LPS and LL-37 challenge, either alone or in combination, for 4 h. IL-8 secretion was quantified using ELISA. Data are shown as means ± SEM (n = 3 independent experiments done in triplicate). *P* < 0.05 (one-way ANOVA *post hoc* Bonferroni correction for multiple group comparison or two-tailed Student’s *t*-test for two groups) was considered significant.

To further ascertain upstream effectors in the p38MAPK pathway, we investigated TLR4 as a main cofactor in the LPS/LL-37 effect. Knocking-down TLR4 (shRNA: Fig 6D) or blocking TLR4 with an antagonist (LPS-RS: Fig 6C) blocked p38MAPK phosphorylation induced by LL-37/LPS. A TLR4 and EGFR interaction through Src kinases in activating p38 MAPK was then assessed. Src kinase LYN allowed cross-talk between TLR4 and EGFR in human mammary epithelial cells, inhibition of which protected mice against LPS induced (i.p.) lethality (45). We observed that Src inhibition (PP1) reduced activation of p38MAPK (Fig 6C), resulting IL-8 in HT29 stimulated by LPS/LL-37 (Fig 6E). Therefore, cathelicidin and LPS increased IL-8 secretion in colonic epithelial cells via p38MAPK activation which may be downstream of TLR4, Src kinases, and EGF receptors.

### MEK1/2 mediated NF-κB activation regulates IL-8 expression in colonic epithelium induced by LL-37/LPS

Activation of NF-κB is essential for IL-8 synthesis in airway and cervical epithelial cells (46, 47). Pharmacological inhibition of the NF-κB activating IKK complex (PS1145) blocked upregulation of *IL-8* mRNA and secretion by HT29 cells stimulated by LL-37/LPS (Figs 7A (i) and (ii), respectively). The ability of LL-37/LPS to stimulate NF-κB promoter elements upstream of IL-8 was further assessed with a luciferase reporter construct (48); LPS/LL-37 induced NF-κB promoter in HT29 cells. Luciferase reporter activity was abolished upon mutation at the NF-κB promoter site, whereas it remained after mutations at AP-1 or NF-IL6 sites (Fig 7B). Likewise, LPS/LL-37 induced time-dependent phosphorylation of the p65-NF-κB subunit (Fig 7C). This LPS/LL-37 mediated NF-κB p65 subunit phosphorylation was partially downregulated in TLR4 knock-down cells (Fig 7D). NF-κB activation is regulated by MEK1/2 in LPS-stimulated human renal and alveolar epithelial cells (49, 50). Similarly, we found inhibition of MEK1/2 blocked phosphorylation-transactivation of the NF-κB subunit p65 (Fig 7E (i)) in HT29 cells exposed to LL-37/LPS, together with reduced IL-8 secretion (Fig 7E (ii)). Inhibition of p38MAPK, Src, and EGFR kinase had no effect on LL-37/LPS induced phosphorylation of NF-κB/p65 (Fig 7F). Therefore, upregulation of *IL-8* mRNA and its protein translation were contigent on two pathways: one NF-κB dependent and downstream of TLR4 and MEK1/2 MAPK and another NF-κB independent, regulated by Src, the EGFR kinase, and p38MAPK.

**Fig 7.**
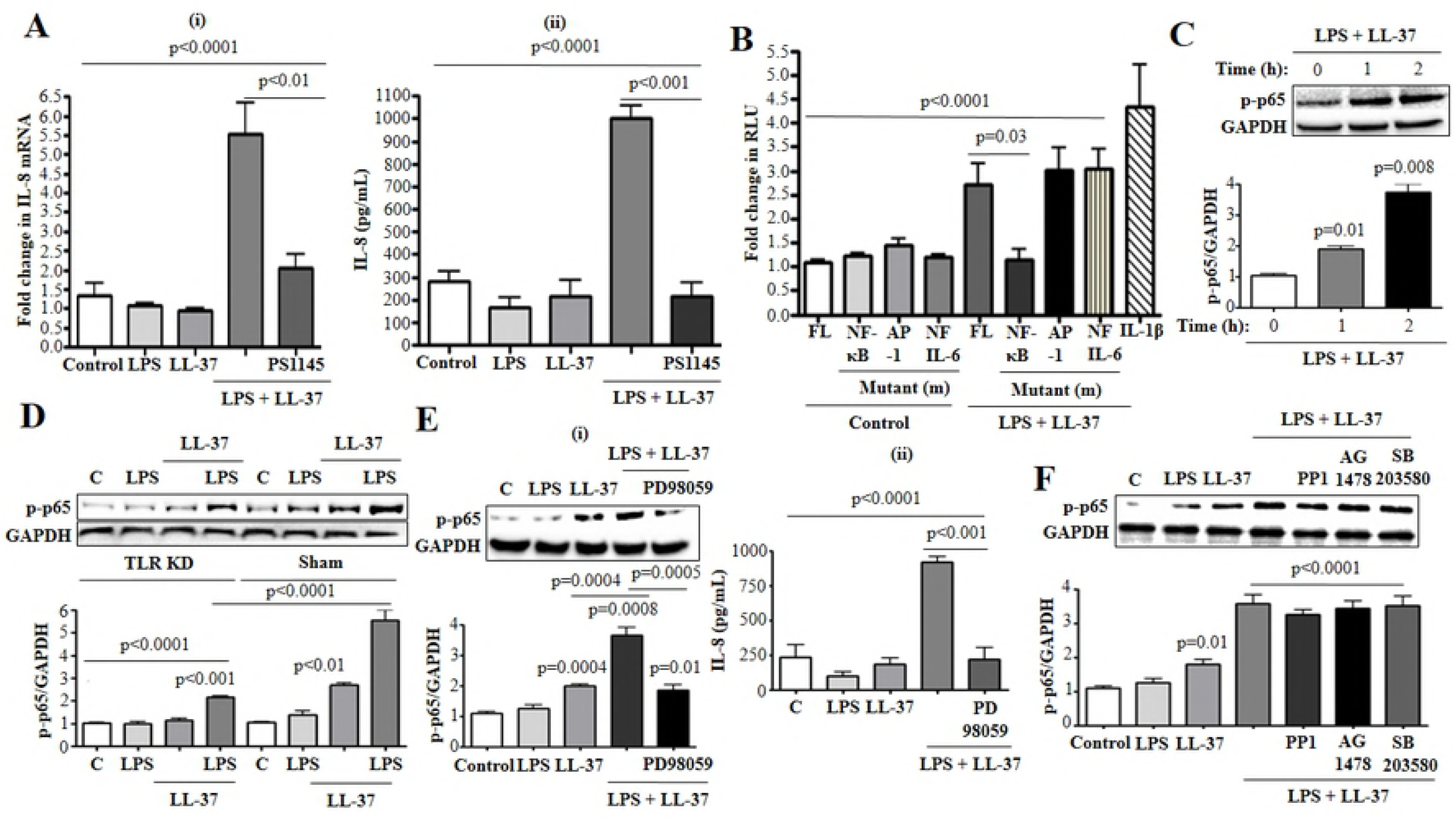
NF-κB activation is needed for IL-8 synthesis induced by cathelicidin and LPS in colonic epithelial cells. (**A *i-ii***) HT29 cells were pre-treated with NF-κB inhibitor PS1145 (1 μM) for 1 h, followed by treatment with LPS (1 μg/mL) and LL-37 (10 μg/mL). (**A *i***) *IL-8* gene synthesis was quantified after 1 h using qPCR (n= 4) and, (**A *ii***) protein secretion was assessed after 4 h using ELISA. **(B)** HT29 cells were transfected with 174bp wild type (FL) or mutant IL-8 promoter construct (containing individual mutated sites for either NF-κB or AP-1 or NF-IL-6) upstream of luciferase gene, followed by stimulation with LPS and LL-37 for 4 h. Data are represented as fold change in relative light units (RLU), normalized to renilla luciferase transfection control. IL-1β (10 ng/mL) was used as positive control. **(C)** LPS-LL-37 mediated time dependent activation of NF-κB in HT29 cells was confirmed using western blotting against phosphorylated p65 subunit of NF-κB. **(D-F)** HT29 cells were either **(D)** sham transfected/knocked down for TLR4 (TLR4 KD), or (E and F) treated with inhibitors for **(E)** MEK1/2 (F) EGFR kinase, Src kinases, p38MAPK, denominated as PD98059 (20 μM), AG1478 (1 μM), PP1 (1 μM) or SB203580 (2 μM), respectively, for 1 h, followed by LPS/LL-37 treatment, either alone or in combination for 2 h (**D, E *i* and F**) or 4 h (**E *ii***). p65 phosphorylation was assessed by western blotting with specific antibodies. Total loading was confirmed after blotting for GAPDH. (**E *ii***) IL-8 secretion was quantified using ELISA. Data are shown as means ± SEM (n = 3 independent experiments done in triplicate). *P* < 0.05 (one-way ANOVA *post hoc* Bonferroni correction for multiple group comparison or two-tailed Student’s *t*-test for two groups) was considered significant.

### p38MAPK/MEK1/2-mediated IL-8 mRNA stabilization contributes to IL-8 production in colonic epithelium stimulated by LPS/LL-37

LPS/LL-37 increased production of IL-8 (Fig 2), although EGFR kinase was mostly involved in post-transcriptional effects (Fig 5). EGFR kinase promotes mRNA stabilization of growth factor amphiregulin in human keratinocytes (HaCaT) upon exposure to ultraviolet B radiation (51). Moreover, downstream effectors p38MAPK and MEK1/2 increase the half life of *IL-8* gene in colonic epithelial cells (52). Thus, we assessed whether LPS/LL-37 modulates IL-8 mRNA half-life. After LPS/LL-37 stimulation for 2 h, actinomycin D was added (which blocked further transcription) either alone or together with a p38MAPK inhibitor (SB203580) or an MEK1/2 inhibitor (PD98059). With actinomycin D alone, *IL-8* mRNA was stable at 1 h, declined at 2 h (~40% of control level), and stabilized (~25%) over 3-4 h (Fig 8). In contrast, inhibition of p38MAPK accelerated the decline in *IL-8* transcripts within 30 min (~60% of control level) and 1 h (~40%) (Fig 8). In the presence of the MEK1/2 inhibitor, there was no decline in *IL-8* transcript at 30 min, but some reduction (~25%) at 1 h relative to the actinomycin D. Thus, LPS/LL-37 via p38MAPK and MEK1/2 signalling could stabilize *IL-8* mRNA, which in turn increased IL-8 production.

**Fig 8.**
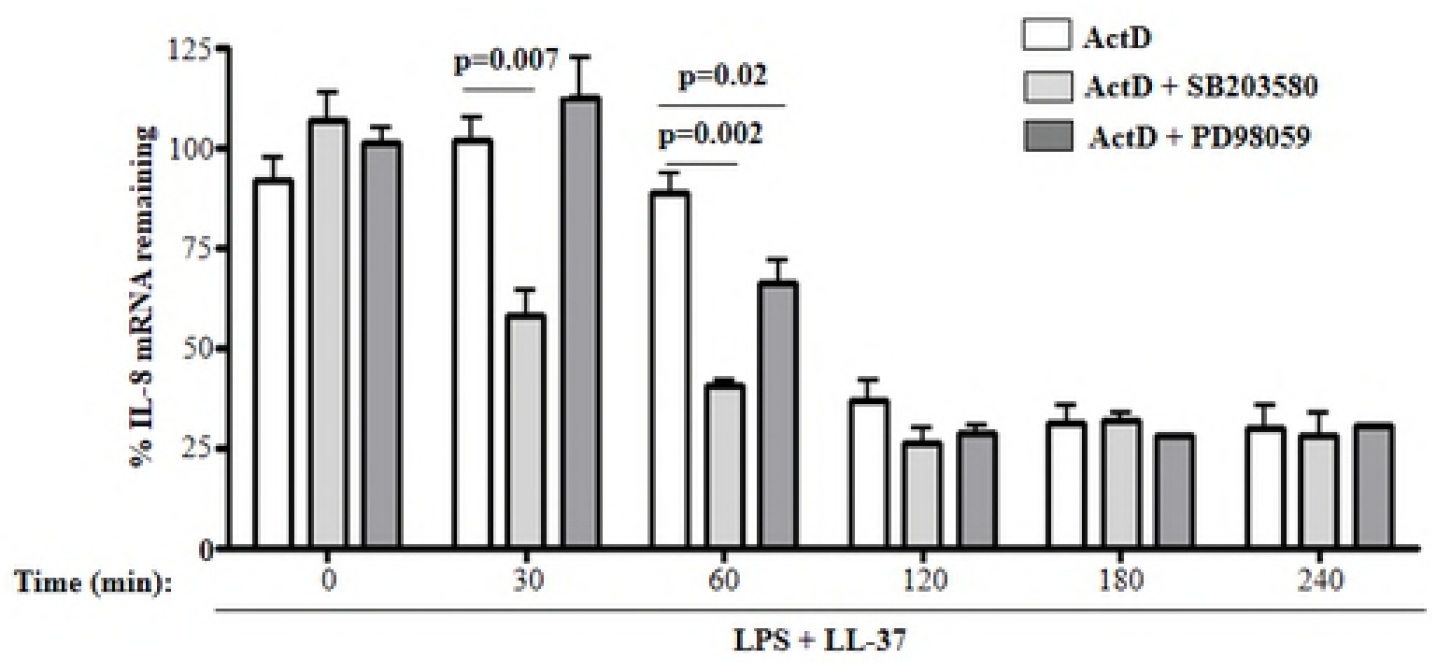
LL-37-LPS synergism promotes p38MAPK and MEK1/2 mediated transcript stabilization in colonic epithelial cells. HT29 cells were stimulated with LPS (1 μg/mL) and LL-37 (10 μg/mL) for 2 h, washed, and incubated with a gene synthesis inhibitor actinomycin D (ActD 1 μM) ± a p38MAPK inhibitor SB203580 (2 μM) or a MEK1/2 inhibitor PD98059 (20 μM) for different time periods (n= 4). Total RNA was extracted and quantified using qPCR. Data are shown as means ± SEM (*n* = 3 independent experiments done in triplicate, unless mentioned otherwise in respective sub-figure). *P* < 0.05 (two-tailed Student’s *t*-test for two groups) was considered significant.

### Secreted colonic IL-8 induced by cathelicidin and LPS promote activation and recruitment of neutrophils in the colon

IL-8 is a potent chemokine acting via CXCR1/2 receptors expressed on the surface of neutrophils (53). Migration of neutrophils into inflamed tissue is essential for local innate immune responses (54, 55). To determined the role of naturally occurring cathelicidins in colonic defenses, C57BL/6 wild type and *Cramp^−/−^* mice were orally challenged with *C. rodentium* and neutrophil infiltration in colons assessed at the peak of infection, day 7 pi (56) (Fig 9A). Wild type mice infected with *C. rodentium* had acute colitis with sloughing of epithelial cells, decreased colon length, and more pronounced colonic hyperplasia compared to *Cramp^−/−^* mice infected with *C. rodentium* (Figs 9B (i-iii)). *Cramp^−/−^* mice during *C. rodentium* had lesser neutrophil infiltration compared to wild type mice, as determined using immunofluorescent Ly6G antibody (Fig 9C) and myeloperoxidase (MPO) activity in the distal colon (Fig 9D). Indeed, neutrophil recruitment and colon MPO activity in *Cramp^−/−^* mice (either sham and *C. rodentium* infected) were similar to that of PBS-treated wild-type mice. KC cytokine production was reduced in colon tissue extracts from *C. rodentium* infected *Cramp^−/−^* mice compared with wild-type infected mice (Fig 9E). Fecal shedding of *C. rodentium* was ~10-fold greater in *C. rodentium* infected *Cramp^−/−^* mice compared with infected wild-type mice (Fig 9F). Taken together, endogenous cathelicidin, perphaps by inducing chemoattractant KC, was necessary for the influx of neutrophils into the colon and *C. rodentium* clearance.

**Fig 9.**
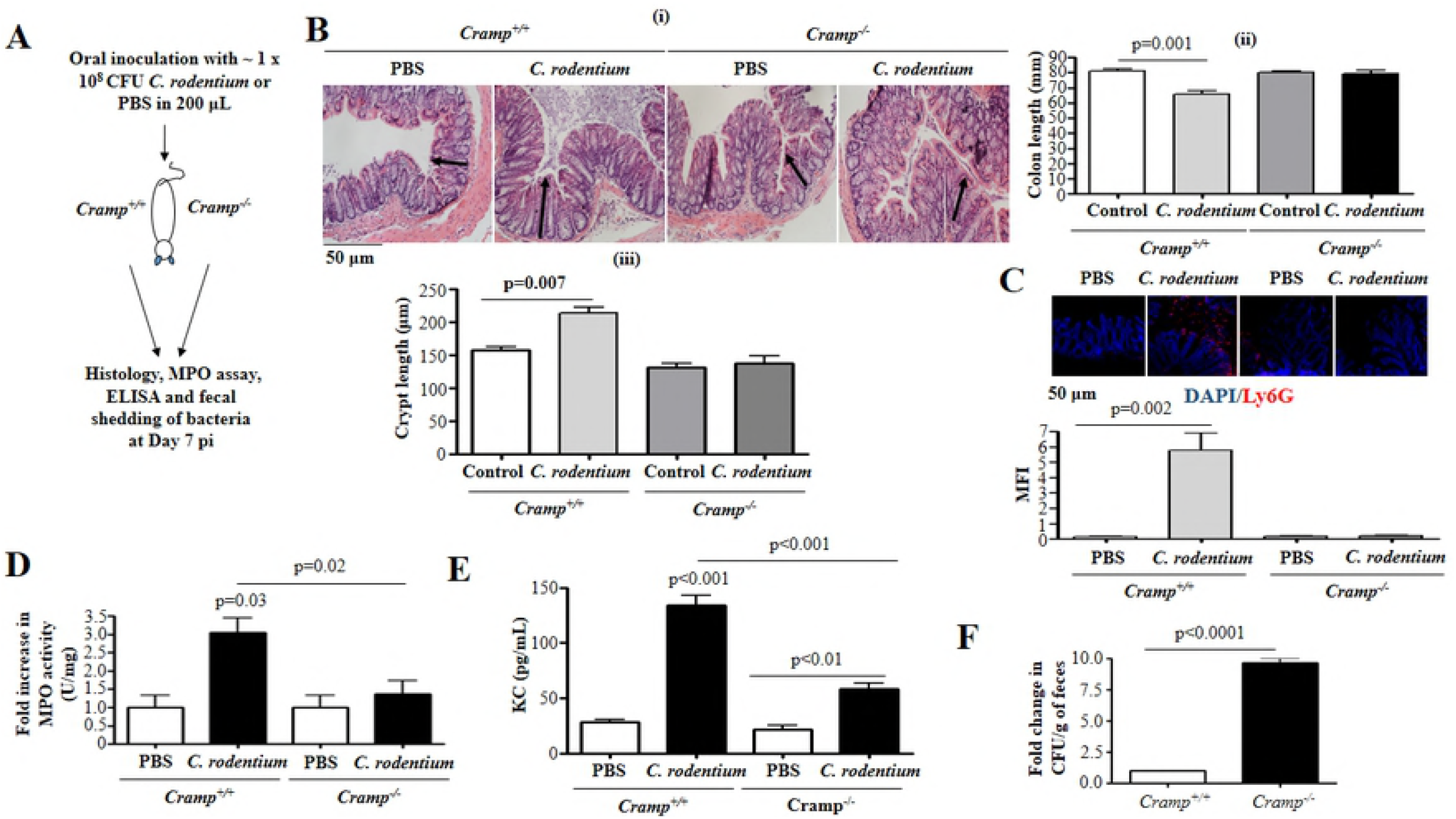
Neutrophil activation in C57BL/6 wild type and Cramp^−/−^ mice challenged with *C. rodentium.* **(A-D)** C57BL/6 *Cramp^+/+^* and *Cramp^−/−^* mice were orally infected with *C. rodentium* (1 x 10^8^ CFU in 200 μL of PBS) for 7 d. **(A)** Experimental design depicting *C. rodentium* infection of mice. **(B *i*)** Hematoxylin-eosin (H&E) staining of mice colonic tissues. Red arrows denote crypt length. (**B *ii-iii***) Histograms depicting colon length (**B *ii***) or crypt length (**B *iii***) in mice at 20X magnification (only for **B *iii***). **(C)** Immunofluorescence staining of mouse colonic tissue with anti-Ly6G antibody (5 μg/mL). Fluorescence was calculated using ImageJ 1.50i software and represented as mean fluorescence intensity (MFI). (**B and C**) Images are representative of n=4 mice. (**D and E**) Bar graphs of MPO activity (D) and KC protein expression **(E)** in murine colonic mucosa after 7 d pi with *C. rodentium.* **(F)** Bar graph of fold change in fecal shedding of *C. rodentium* per gram of feces at 7 d pi. Data are shown as means ± SEM (n=4/group). *P* < 0.05 (One-way ANOVA *post hoc* Bonferroni correction for multiple group comparison or two-tailed Student’s *t*-test for two groups) was considered significant.

## Discussion

Colonic epithelial cells are expected to prevent harmful reactions against the commensal microbiome, but concurrently need to actively recognize invading pathogenic microbes and microbe-associated LPS to initate inflammatory responses. Beyond direct antimicrobial effects of cathelicidins (observed at higher concentrations; ~ 50 μg/mL for *S. typhimurium* (57) than physiologically available at mucosal surfaces of adults (< 20 μg/mL) (58)), this study established that human cathelicidin (LL-37) acted synergistically with *S. typhimurium* or its LPS in intestinal epithelial cells to induce IL-8 biosynthesis. Further, endogenous cathelicidin promoted synthesis of neutrophil KC cytokine in the colon and was necessary for neutrophil inflammatory responses and control of intestinal related Gram negative pathogen *(C. rodentium)*. This role of cathelicidin in the gut epithelium was novel and differed from actions reported in macrophages (59) and corneal fibroblasts (60) in which cathelicidin restricted LPS pro-inflammatory effects.

Mechanistically, we deciphered a coordinated recognition and internalization of LPS/LL-37 complex and detailed signalling pathways for production of IL-8 (Fig 10). First, a physical interaction between cathelicidin and LPS appeared to occur extracellularly, as pre-incubation of polymyxin B with LPS blocked LPS/LL-37 complex-induced production of IL-8. Cathelicidin may either bind to the same site on LPS at which polymyxin B binds, or the peptide triggers conformational changes in LPS structure that mask polymyxin B binding sites. It was noticed that LL-37 sequence has some homology with the B-subunit of cholera toxin, which interacts with membrane GM1 to facilitate internalization of the enzymatically active toxin A-subunit (35). Thus, we propose that, like cholera toxin, internalization of LPS in intestinal epithelial cells is mediated by LL-37 in conjunction with LPS, utilizing a GM1-mediated lipid raft mechanism. In agreement, LL-37 promoted endocytosis of Alexa 488-conjugated-LPS via lipid rafts in lung fibroblast (61) and epithelial cells (17). Of note is that intact LPS, and not simply the lipid A moiety, was needed for LPS/LL-37-mediated IL-8 secretion by HT29 cells. A lack of ability of lipid A on its own to trigger a host response has been reported for *Caenorhabditis elegans* in which an *S. enterica* mutant for LPS, but not a mutant for lipid A, failed to activate MAPK signalling and subsequent gonadal programmed cell death (62). In our study, LPS alone in colonic epithelial cells did not induce IL-8 mRNA transcription or protein translation (for up to 4 h). A temporal regulation of IL-8 synthesis by LPS in colonic epithelium seems to occur; basal IL-8 production was detected after 12 h (63) whereas siginificant increase in secretion above baseline was evident only at 18 h post LPS challenge (up to 1 μg/mL) in HT29 cells (64).

**Fig 10.**
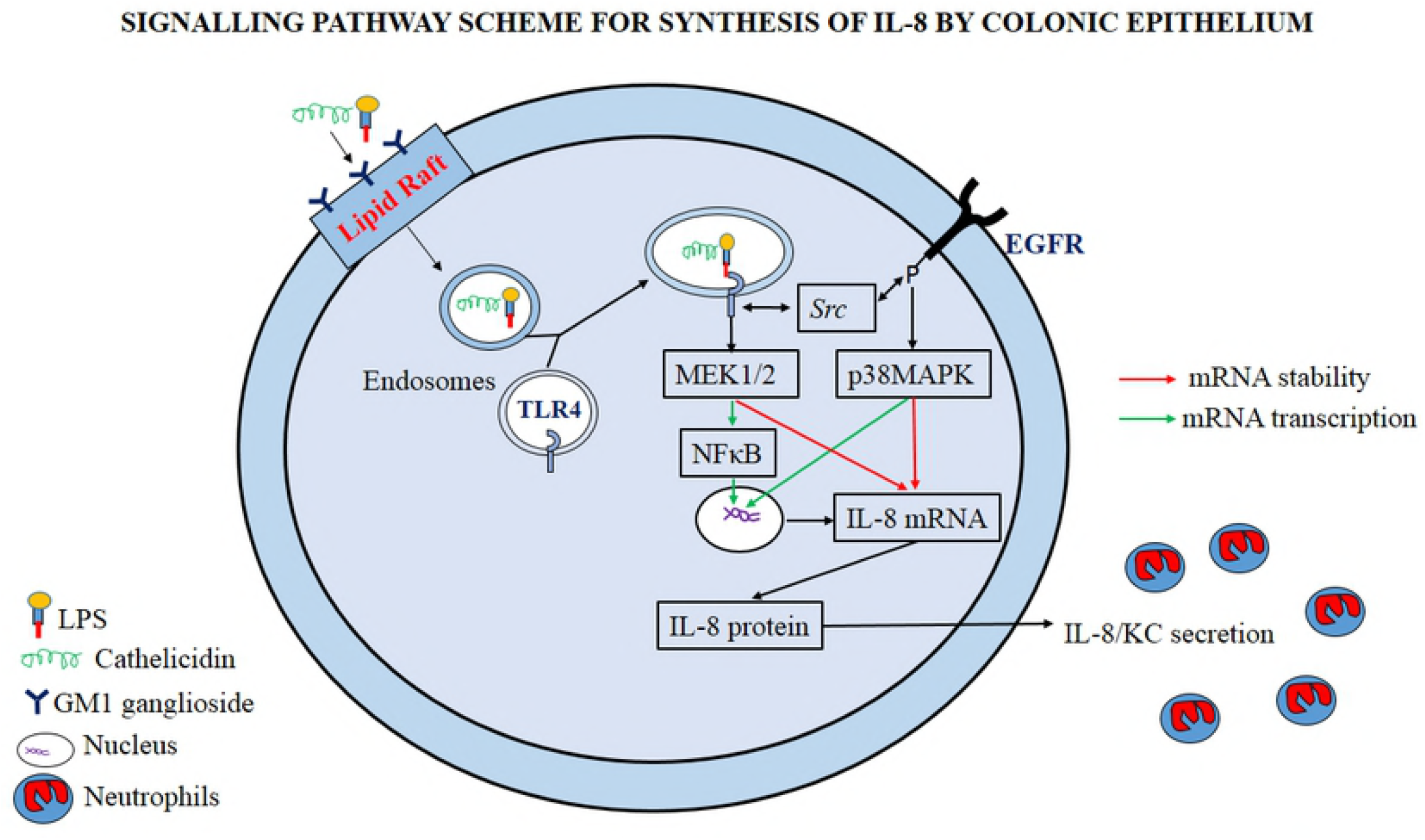
A scheme of the signalling mechanisms elicited by cathelicidin in synergy with LPS to promote IL-8 secretion in colonic epithelium and subsequent neutrophil recruitment/activation. In colonic epithelial cells, LL-37 physically binds and faclilitates LPS uptake via interaction with GM1 (monosialotetrahexosylganglioside) in lipid rafts. Intracellularly, LPS interacts with TLR4 to promote activation of two signalling axes: one NF-κB dependent, which signals via TLR4/MEK1/2, and another NF-κB independent, which relies on TLR4, Src kinase, and EGFR kinase cross-talk and subsequent p38MAPK activation. Whereas NF-κB dependent signalling primarily regulates IL-8 mRNA synthesis, NF-κB independent signalling primarily controls IL-8 mRNA stability. Both signalling pathways together promote colonic IL-8 protein synthesis and secretion. The induced colonic IL-8 chemokine regulates intestinal defenses via neutrophil recruitment/activation.

A LPS/LL-37 complex co-opted via lipid rafts and triggered intracellular TLR4 signalling. A key role of TLR4 within the cell, rather than on the plasma membrane, has been postulated for the mature colonic epithelium to minimally respond to commensal bacteria (65). Lipid rafts were also necessary for LPS signalling via intracellular TLR4 in mouse small intestinal crypt (m-IC_cl2_) cells (66). Moreover, internalization of LPS and interaction with intracellular TLR4 was reported in synthesis of IL-8 in human colonic epithelial cells and human bronchial epithelial cells (17, 63). Besides intracellular TLR4, cathelicidin/LPS induced IL-8 synthesis in colonic epithelium was dependent on enhanced EGFR kinase and Src kinase activity. An integration among TLR4 and EGFR through LYN (Src kinase) regulated plasma concentrations of IL-6, TNF-α and KC in mice systemically challenged with LPS (45).

Synthesis of IL-8 induced by LPS/LL-37 involved MEK1/2 and p38MAPK activation. Unexpectedly, direct inhibition of ERK using specific FR180204 inhibitor (unlike PD98059 which inhibits MEK1/2 to block downstream ERK1/2 activation) (67) did not affect IL-8 either transcription or translation. Similarly, mutation in AP-1 promoter site, an ERK target (68), did not affect luciferase-tagged IL-8 promoter construct activity in response to LPS/LL-37 (Fig 7B). Perhaps a non-canonical MEK1/2 signalling (independent of ERK activation) is involved in colonic IL-8 synthesis induced by LPS/LL-37 complex. In particular, we determined that MEK1/2 was directly involved in NF-κB (p65) phosphorylation/activation in colonic epithelium. Such alternative MEK1/2 signalling was reported in intestinal epithelial (T84) cells stimulated in proliferation/survival by fractalkine (CX3CL1, a chemoattractant for T cells and monocytes) (69, 70), although IL-8 production was not assessed. In regards to p38MAPK, LPS/LL-37 induced activation/phosphorylation of p38MAPK in colonic cells was partially dependent on EGFR and Src kinase activity. Such a signalling axis (EGFR-Src-p38MAPK) has been demonstrated in EGF-dependent young adult mouse colon (YAMC) cell proliferation/migration and hence, closure of mechanically-induced wounds (71). Conversely, p38MAPK was not directly involved in NF-κB (p65) phosphorylation at Ser^536^ (p-Ser 536) site. Perhaps, p38MAPK regulated IL-8 transcriptional activity through phosphorylation of NF-κB (p65) at Ser^276^, as demonstrated in RAW264.7 macrophage-like cells (72).

On another level of IL-8 regulation, signalling via p38MAPK and MEK1/2 kinases has an impact on IL-8 mRNA turnover, prolonging mRNA transcript lifetime. Likewise, p38MAPK and MEK1/2 extended IL-8 mRNA half-life in HT-29 cells stimulated with TNF-α (52). Stabilization of IL-8 mRNA is crucial for a sustained increase in IL-8 secretion, as *IL-8* gene has a short half-life (~ 24 min), as determined in untreated THP-1 monocytes using an actinomycin D chase assay (73). Therefore, colonic epithelial cells may use a specific combination of signalling pathways (e.g. TLR4, Src/EGFR/MAP kinases and NF-κB) to regulate IL-8 gene transcription, mRNA stabilization and translation in response to LPS/LL-37. The proposed combined pathways (Fig 10), one NF-κB dependent and downstream of TLR4 and MEK1/2 MAPK and another NF-κB independent, regulated by Src, the EGFR kinase, and p38MAPK, both contribute to the overall understanding of IL-8 synthesis by the colonic epithelium.

The IL-8 synthesis by colonic epithelium when exposed to *S. typhimurium*/LPS and stimulated with cathelicidin was biologically relevant, stimulating calcium signalling and neutrophil elastase secretion in human blood-derived neutrophils. Thus, colonic IL-8 may indirectly enhance killing roles in recruited neutrophils that are essential for *S. typhimurium* clearance (55). The function of KC/LL-37 to modulate *C. rodentium* induced colitis reinforced, in two ways, the observed action of LPS/LL-37 to induce IL-8 in colonic epithelium. First, lack of presumed LPS/cathelicidin induction of KC resulted in reduced infiltration of neutrophils in cathelicidin-null mice at day 7 pi (Figs 9C-D). In agreement, there was reduced colitis in cathelicidin-deficient mice, relative to the wild-type mice, at day 14 pi (74). In addition, *C. rodentium* infected cathelicidin-null mice also had lesser signs of colitis, with reduced crypt hyperplasia and colon shortening (Figs 9B (ii-iii)). Second, elimination of *C. rodentium* was markedly impaired in cathelicidin-deficient mice. This role of cathelicidin to promote clearance of *C. rodentium* has been reported in *Cramp^−/−^* mice, which had higher systemic dissemination than *Cramp^+/+^* mice (74). Based on our work, defective clearance of the infection may be attributed to the reduced KC-stimulated influx of inflammatory cells (i.e., neutrophils) in cathelicidin-null mice. Thus, cathelicidin has a dual role, contributing both to the neutrophil attack and elimination of invading pathogens. Such immunomodulatory functions of cathelicidin may be more relevant in gut defense than direct killing of microbes, as lower peptide concentrations are required for regulating cytokine production *in vitro (75).* However, although we confirmed KC/cathelicidin contributed to the neutrophil influx and bacterial clearance in *C. rodentium* colitis, cathelicidin-null mice had more severe colitis induced by dextran sodium sulphate (DSS) (76). Thus, the role for cathelicidin in intestinal pathophysiology may depend on the noxious agent and be specific to the inflammatory milleu.

In conclusion, this study demonstrates that cathelicidin can act as a sensor of either *S. typhimurium* or LPS to form an LPS-LL-37 complex, internalized via GM1-lipid rafts, and stimulates intracellular TLR4-mediated upregulation of IL-8. The IL-8 secreted by colonic epithelium directs the influx of neutrophils into the colon and pathogen clearance during colitis. Proactive stimulation of colonic IL-8 by cathelicidin may complement its direct antimicrobial activity. These uncovered immunomodulatory roles of cathelicidin highlight the therapeutic potential of endogenous and supplemented exogenous cathelicidins for gut innate defense and the clearance of intestinal pathogens.

## Acknowledgements

The authors thank D Proud (University of Calgary) for assisting in the experimental designs and providing plasmid constructs for IL-8 and J Kastelic (University of Calgary) for editing this manuscript. Immunofluorescence studies were conducted in the Live Cell Imaging Facility, Snyder Institute, University of Calgary.

## Supporting information

**S1 Fig. Time dependent development of colonoids from human biopsies and mouse colonic tissue.** Mice and human iPSCs were isolated and grown on 12-well plates. The figure depicts colonic tissue acquiring colonoid architecture with detectable lumen and developing crypts at Day 10 post culturing.

**S2 Fig. Dose curve for IL-8 secretion in colonic epithelium in presence of either LPS or LL-37 alone.** HT29 cells were stimulated with variable doses of **(A)** LPS and **(B)** LL-37 alone for 4 h, followed by quantification of IL-8 protein secretion (pg/mL) using ELISA. Data are shown as means ± SEM (*n* = 3 independent experiments done in triplicate). *P* < 0.05 (One-way ANOVA *post hoc* Bonferroni correction) was considered significant.

**S3 Fig. Secreted IL-8 from colonic epithelium stimulated by cathelicidins and LPS induces calcium flux and activation of human primary neutrophils. (A-B)** Human neutrophils were seeded at **(A)** 1 x 10^6^ cells and pre-incubated with Fluo4-NW dye (45 m, RT) or **(B)** 1 x 10^4^/well in an 8-well chamber and incubated at 37° for 1 h. (A-B) Cells were then exposed to supernatant from HT29 cells unstimulated (control) or stimulated with LPS and LL-37, either alone or in combination, for 4 h ± anti-IL-8 antibody (1 μg/mL) or CXCR1/2 inhibitor SCH527123 (20 μM). rIL-8 was used as positive control. IgG was used as an isotype control. **(A)** Data are represented as percentage fluorescence emission at 530 nm of positive control calcium ionophore A23187 (CI A23187). **(B)** Neutrophil activation was assessed by neutrophil elastase secretion (depicted by red arrows) using immunocytochemistry using anti-neutrophil elastase antibody (5 μg/mL). Data are represented as fold increase in MFI normalized to respective control, for three independent experiments. Data are shown as means ± SEM (*n* = 3 independent experiments done in triplicate). *P* < 0.05 (one-way ANOVA *post hoc* Bonferroni correction for multiple group comparison or two-tailed Student’s *t*-test for two groups) was considered significant.

**S4 Fig. RasMol analysis of cholera toxin B subunit (CTxB) and active form of human cathelicidin LL-37.** Areas of structural similarity between CTxB and LL-37 using webPRANK software from EMBL-EBI (http://www.ebi.ac.uk/goldman-srv/webprank/). The reliability index (confidence in structural similarity at a particular pair of alignment) is graded as shades of grey, with black representing maximum reliability.

**S5 Fig. LPS and LL-37 promotes p38MAPK phosphorylation in colonic epithelium cells.** HT29 cells were stimulated with LPS (1 μg/mL) and LL-37 (10 μg/mL) for 4 h. The figure is a representative image of three independent experiments.

